# Phylogenetic analyses of AGO/DCL/RDR proteins in green plants refine the evolution of small RNA pathways

**DOI:** 10.1101/2022.01.18.476847

**Authors:** Sébastien Bélanger, Junpeng Zhan, Blake C. Meyers

**Affiliations:** Donald Danforth Plant Science Center, St. Louis, MO, USA, 63132; Division of Plant Science and Technology, University of Missouri, Columbia, MO, USA, 65211

**Keywords:** Small RNA, Viridiplantae, AGO, DCL, RDR, DRB, SE, SGS3, HEN1, evolution

## Abstract

Several protein families play a role in the biogenesis and function of small RNAs (sRNAs) in plants. Those with primary roles include Dicer-like (DCL), RNA-directed RNA polymerase (RDR), and Argonaute (AGO) proteins. Protein families such as double-stranded RNA-binding (DRB), SERRATE (SE), and SUPPRESSION OF SILENCING 3 (SGS3) act as partners of DCL or RDR proteins. Here, we present curated annotations and phylogenetic analyses of seven sRNA pathway protein families performed on 196 species in the Viridiplantae (aka green plants) lineage. Our results suggest that the RDR3 proteins emerged earlier than RDR1/2/6. RDR6 is found in filamentous green algae and all land plants, suggesting that the evolution of RDR6 proteins coincides with the evolution of phased small interfering RNAs (siRNAs). We traced the origin of the 24-nt reproductive phased siRNA-associated DCL5 protein back to *Acorus americanus*, the earliest diverged, extant monocot species. Our analyses of AGOs identified multiple duplication events of *AGO* genes that were lost, retained or further duplicated in sub-groups, indicating that the evolution of *AGOs* is complex in monocots. The results also refine the evolution of several clades of AGO proteins, such as AGO4, AGO6, AGO17 and AGO18. Analyses of nuclear localization signal sequences and catalytic triads of AGO proteins provide insights into the regulatory roles of diverse AGOs. Collectively, this work generates a curated and evolutionarily coherent annotation for gene families involved in plant sRNA biogenesis/function and provides insights into the evolution of major sRNA pathways.

## INTRODUCTION

Small RNAs (sRNAs), non-coding RNAs that are usually 20 to 24 nucleotides (nt) in length, are key regulators of plant development, genome integrity, and environmental response (Borges and Martienssen, 2015). Plant sRNAs are primarily classified as microRNAs (miRNAs) and small interfering RNAs (siRNAs) (Axtell, 2013). miRNAs originate from single-stranded transcripts (i.e. pri-miRNAs) produced by RNA polymerase II (Pol II). The pri-miRNAs fold into hairpin-like structures that are cleaved by a ribonuclease III enzyme called DICER-LIKE 1 (DCL1) to produce miRNAs, which are ∼21 nt in length (Borges and Martienssen, 2015). miRNAs mainly function via guiding transcript cleavage by Argonaute (AGO) proteins or mediating translational repression (Borges and Martienssen, 2015).

siRNAs can be sub-classified into heterochromatic siRNAs (hc-siRNAs) and secondary siRNAs (Axtell, 2013); the latter includes phased secondary siRNAs (phasiRNAs) and epigenetically activated siRNAs (easiRNAs). hc-siRNAs are 24 nt in length and their precursors are produced from heterochromatic genomic regions by Pol IV (Axtell and Meyers, 2018). PhasiRNAs originate from protein-coding transcripts or long non-coding RNAs, which are Pol II transcripts (Fei et al., 2013; Liu et al., 2020). PhasiRNA biogenesis initiates via miRNA-directed, AGO-catalyzed cleavage of a single-stranded RNA precursor, which is then converted to a double-stranded RNA (dsRNA) by an RNA-dependent RNA polymerase (RDR), and processed into 21-nt or 24-nt RNA duplexes by a DCL protein (Fei et al., 2013; Liu et al., 2020). PhasiRNAs can be further sub-classified as those derived from protein-coding loci and those derived from long non-coding loci (Liu et al., 2020). EasiRNAs are 21- or 22-nt in length and enriched in pollen vegetative cells (Creasey et al., 2014). hc-siRNAs function mainly in transposon silencing by mediating DNA methylation (i.e. RNA-directed DNA methylation) (Matzke and Mosher, 2014; Borges and Martienssen, 2015; Axtell and Meyers, 2018). 21-nt phasiRNAs, including *trans*-acting siRNAs (tasiRNAs) and 21-nt reproductive phasiRNAs, mediate cleavage of target transcripts (Fei et al., 2013; Tamim et al., 2018; Jiang et al., 2020; Zhang et al., 2020), while 24-nt tasiRNAs and 24-nt reproductive siRNAs have been implicated in mediating DNA methylation in *cis (Wu et al., 2012; Zhang et al., 2021)*. The diversity of molecular functions of plant sRNAs are likely associated with the involvement of specific biogenesis (e.g., DCL and RDR) or effector (e.g., AGO) proteins.

Four *DCL* genes are present in the Arabidopsis (*Arabidopsis thaliana*) genome and are named *DCL1* to *DCL4* (Voinnet, 2009). DCL1 produces miRNAs from pri-miRNAs (Kurihara and Watanabe, 2004; Hiraguri et al., 2005); DCL2 produces viral-derived and endogenous 22-nt siRNAs (Gasciolli et al., 2005; Ding and Voinnet, 2007; Parent et al., 2015; Taochy et al., 2017; Wu et al., 2017; Wang et al., 2018; Jia et al., 2020); DCL3 produces hc-siRNAs, which are 24 nt in length, mainly from Pol IV-derived short transcripts (Blevins et al., 2015; Zhai et al., 2015a; Wang et al., 2021a); and DCL4 processes 21-nt phasiRNAs, including tasiRNAs and 21-nt reproductive phasiRNAs (Gasciolli et al., 2005; Liu et al., 2020) (Table 1). A fifth DCL protein, DCL5 (formerly DCL3b) emerged and evolved in monocots, with a specialized role in producing 24-nt reproductive phasiRNAs in anthers (Song et al., 2012; Teng et al., 2020). The *AGO* gene family is highly diversified in angiosperms, forming three major phylogenetic clades, referred to as AGO1/5/10, AGO2/3/7, and AGO4/6/8/9 (Vaucheret, 2008; Zhang et al., 2015). The selective loading of sRNAs by AGO proteins enables fine regulation of gene expression in plants. In Arabidopsis, sRNA sorting and loading onto AGO are determined or influenced by the 5’ nucleotide and length of sRNA (Bologna and Voinnet, 2014). For example, AGO1 is a canonical effector of miRNAs with a 5’-U, and AGO4/6/9 load hc-siRNAs (Table 1). Whether an sRNA mediates RNA cleavage is determined by the presence or absence of a triad of amino acids (i.e. catalytic triad) in the P-element-induced wimpy testis (PIWI) domain of AGO proteins (Carbonell et al., 2012). In Arabidopsis, AGO1, AGO2, AGO4, AGO7, and AGO10 show slicer activity (Carbonell et al., 2012). The Arabidopsis genome encodes six RDRs (Bologna and Voinnet, 2014), with diverged roles in siRNA biogenesis: RDR1 mainly functions in biogenesis of virus-derived siRNAs (Garcia-Ruiz et al., 2010; Wang et al., 2010; Cao et al., 2014), RDR2 functions in hc-siRNA biogenesis by converting Pol IV transcripts to dsRNAs (Chan et al., 2004; Xie et al., 2004), RDR6 is involved in biogenesis in phasiRNAs (Kumakura et al., 2009; Jouannet et al., 2012) (Table 1), whereas the substrates and functions of RDR3a/b/c (formerly RDR3/4/5 (Wassenegger and Krczal, 2006)) are as-yet largely unknown.

**Table 1.** Features of key AGO, DCL and RDR proteins known to function in the hc- siRNA and phasiRNA pathways.

| Protein | Plant model | Features |  | References |
| --- | --- | --- | --- | --- |
|  |  | sRNA length (nt)<sup>1</sup> | siRNA pathway |  |
| DCL3 | Arabidopsis | 24 | hc-siRNA | (Wang et al., 2021a) |
| DCL4 | Arabidopsis, rice | 21 | phasiRNA | (Xie et al., 2005; Liu et al., 2007; Song et al., 2012) |
| DCL5 | Maize, rice | 24 | phasiRNA | (Song et al., 2012; Teng et al., 2020) |
| RDR2 | Arabidopsis | 24 | hc-siRNA | (Willmann et al., 2011; Hua et al., 2021) |
| RDR6 | Arabidopsis, rice | 24 | phasiRNA | (Peragine et al., 2004; Willmann et al., 2011; Hua et al., 2021) |
| AGO5c | Rice, maize | 21 | phasiRNA | (Komiya et al., 2014; Lee et al., 2021) |
| AGO4 | Arabidopsis | 24 | hc-siRNA | (Rowley et al., 2011) |
| AGO6 | Arabidopsis | 24 | hc-siRNA | (Zheng et al., 2007) |
| AGO18 | Maize, rice | 21 and 24 | phasiRNA | (Sun et al., 2019) |

<sup>1</sup>Typical length of sRNAs associated with these proteins.

As the main biogenesis/effector proteins of sRNAs, the DCL/AGO/RDR proteins often form complexes with other proteins to function. Several members of the DOUBLE-STRAND RNA BINDING (DRB) family interact with DCLs and regulate sRNA biogenesis; for example, in Arabidopsis, DRB1 [aka HYPONASTIC LEAVES1 (HYL1)] protein is a canonical partner of DCL1 in miRNA biogenesis and DRB4 interacts with DCL4 in the biogenesis of 21-nt phasiRNAs (Hiraguri et al., 2005; Nakazawa et al., 2007; Curtin et al., 2008; Fukudome and Fukuhara, 2017). SE is another canonical partner of DCL1 (Machida et al., 2011). In addition, HEN1 interacts with DCL1 to catalyze methylation of miRNAs (Yu et al., 2005; Baranauskė et al., 2015), while also catalyzing methylation of other sRNAs to stabilize them (Yang et al., 2006a). Finally, SGS3 is a canonical protein partner of RDR6 to synthesize dsRNAs (Kumakura et al., 2009). The biogenesis and regulatory functions of sRNAs are often regulated/influenced by these proteins (via interaction with AGO/DCL/RDR proteins), which we refer to collectively as ‘accessory proteins’ of sRNA pathways.

Here, we performed a genome-wide annotation and phylogenetic analyses of the AGO/DCL/RDR/DRB/SE/SGS/HEN1 family proteins in 196 Viridiplantae species to understand the evolution of sRNA pathways in plants, and we examined the nuclear localization signal (NLS) sequences and catalytic triads of AGO proteins. The analyses provide an evolutionarily coherent and standardized annotation for the sRNA biogenesis/effector protein families, refines the evolution of each protein family in green plants, including the ancestral status of RDR3 and the origins of DCL5, AGO17, and AGO18, and provides insights into the evolution of the major sRNA pathways.

## RESULTS AND DISCUSSION

### Annotation of sRNA pathway proteins across ∼200 species in the Viridiplantae lineage

To understand the evolution of sRNA pathways in plants, we annotated and performed a phylogenetic analysis of the RDR, DCL, AGO, SGS3, DRB, SE, and HEN1 protein families in 196 plant genomes, including several plant genomes that were sequenced in the past few years, plus 11 non-plant genomes as outgroups. The analyzed species include four Chlorophyta and 192 Streptophyta species, and 11 outgroup species (including three Rhodophyta, one Oomycota, one Amoebozoa, one fungus and five Metazoa) (Supplemental Table 1). The Streptophyta species included three filamentous green algae, two Marchantiopsida (liverwort), five Bryopsida (mosses) and 182 Tracheophyta species comprising one Lycopodiopsida (lycophyte), three Polypodiopsida (ferns), seven Acrogymnospermae (gymnosperms) and 171 Mesangiospermae (angiosperms). The angiosperms included six basal-most angiosperms (one Amborellales and three Nymphaeales), ten Magnoliids, 58 Liliopsida (monocots) and 99 Pentapetalae (88 core eudicots and 11 basal eudicots including three Proteales, seven Ranunculales and one Trochodendrales) (Figure 1a; Supplemental Data 1; Supplemental Table 1). Together, a total of 2,979 AGO, 1,036 DCL, 1,440 RDR, 2,000 DRB, 470 SE, 455 SGS3, and 224 HEN1 family proteins in the 207 species, including the outgroup species, were annotated and curated for the presence of conserved functional domains (Figure 1b). We built, rooted, and reconciled the protein trees with a species tree inferred from whole proteomes, and inferred the duplication and loss events with an emphasis on monocots, including economically important crop species.

**Figure 1.**
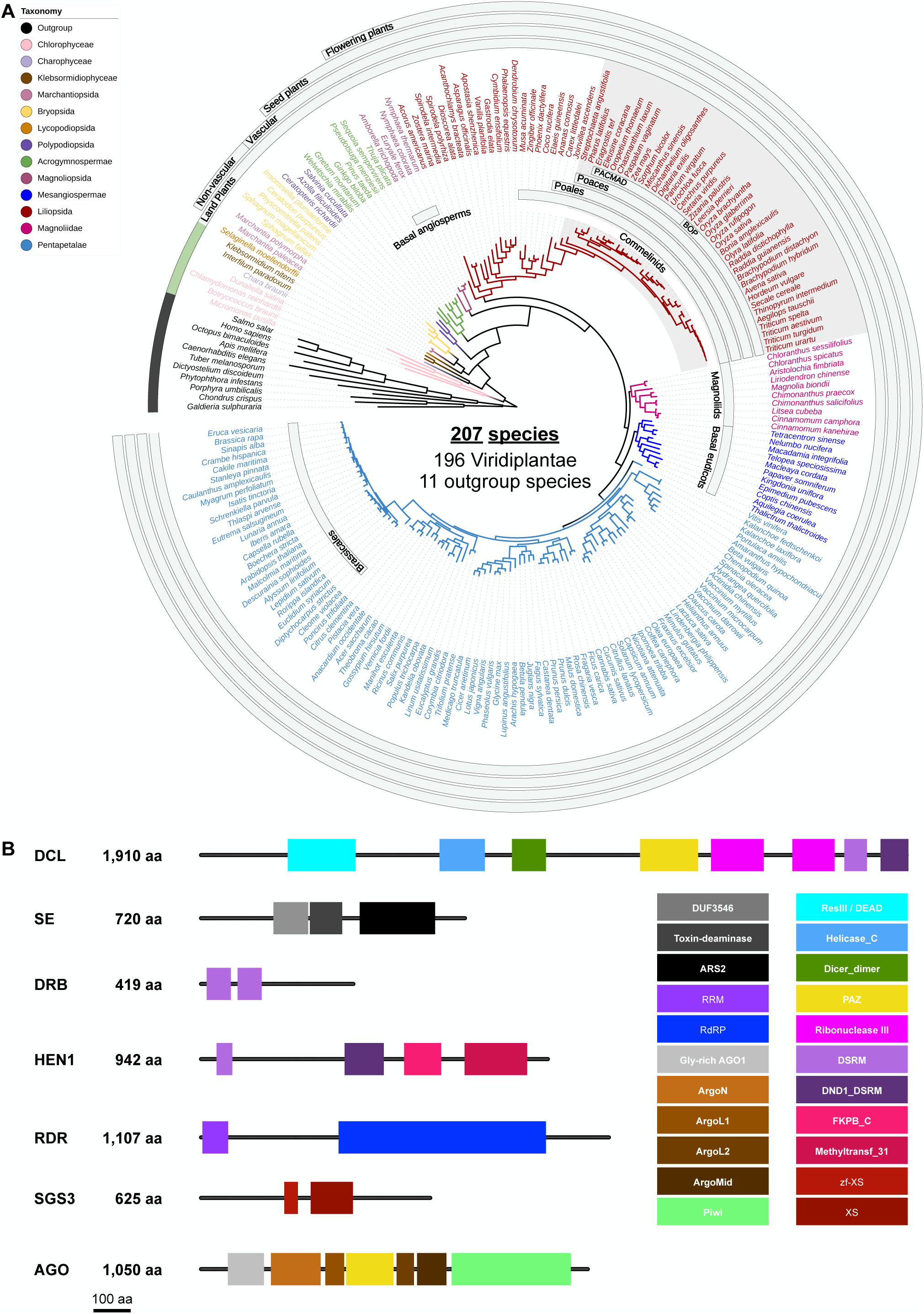
Species and protein families analyzed in this study. **(A)** Species tree of all the examined plant and non-plant species. The tree was generated using OrthoFinder based on whole proteome sequences and visualized using iTOL. **(B)** Canonical domains of each protein family. Presence of these domains were used as the criterion to filter for proteins that are included in the protein trees.

### Molecular evolution of RDRs in Viridiplantae

RDR proteins function in converting single-stranded RNA molecules to dsRNAs, which is a key step in siRNA biogenesis (Borges and Martienssen, 2015). To gain insights into the evolution of siRNA pathways, we performed a phylogenetic analysis of the RDR family proteins. A maximum likelihood tree of 1,440 RDR proteins (including 11 non-plant RDR proteins) was inferred and reconciled with the species tree (Figure 2; Supplemental Data 2; Supplemental Table 2). We detected a total of 13 RDR proteins in four of the 11 outgroup species and two of four Chlorophyta species, including 11 adjacent to the RDR1/2/6 group and two in the RDR3 group, suggesting an early advent of *RDR* genes, likely before the emergence of green plants. Within Viridiplantae, we observed three major monophyletic groups (RDR1/2, RDR3, and RDR6), with RDR6 sister to RDR1 and RDR2, and RDR3 sister to RDR6 and RDR1/RDR2 (i.e., a tree topology of RDR3,(RDR6,(RDR2,RDR1))); this is consistent with previous phylogenetic analyses focused on angiosperms including Arabidopsis (Liu et al., 2014; Sabbione et al., 2019). Furthermore, our results demonstrate that RDR3 is likely ancestral to the other types of RDRs, as an RDR3 protein was found in Chlorophyceae, whereas the other RDRs seem to have emerged in gymnosperms (RDR1 and RDR2) or Klebsormidiophyceae (RDR6).

**Figure 2.**
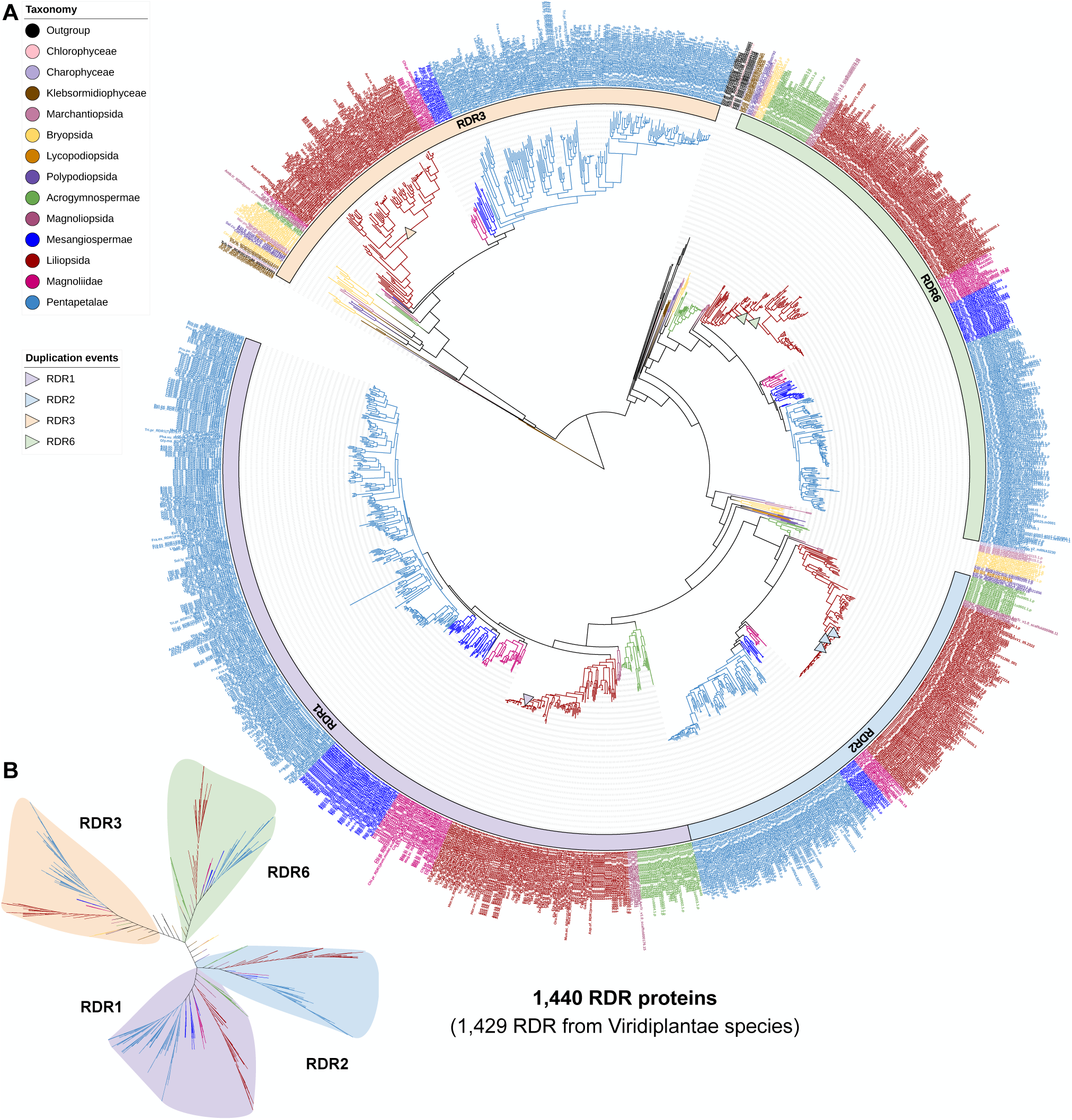
Maximum-likelihood phylogeny of all RDR proteins annotated in the analyzed genomes. **(A)** Rooted phylogeny of all RDR-type proteins in the 207 plant and non-plant species. The particularly numerous nodes marked in royal blue and dark red indicate eudicot and monocot species, respectively. (B) Unrooted view of the major clades of the phylogenetic tree in (A).

The Arabidopsis RDR3a/b/c proteins reside in the same clade, consistent with prior reports (Wassenegger and Krczal, 2006; Hua et al., 2021). For simplicity, we refer to this clade as the RDR3 clade (Supplemental Table 2). Proteins in the RDR3 clade likely emerged at or right before the emergence of green algae, as their orthologs were detected in fungi but not in the Amoebozoa, Oomycota or Metazoa species. Although absent in four of the seven gymnosperms, we found RDR3-clade proteins in all land plant lineages with an evolutionary pattern of (N-S,(G,(Mo,(Ma,E)))) (N-S, non-seed land plants; G, gymnosperms; E, eudicots; Ma, magnoliids; Mo, monocots) (Figure 2). The functions of RDR3-clade proteins remain poorly characterized with incomplete data from Arabidopsis and only overexpression data (which are prone to artifacts and indirect effects) in rice (*Oryza sativa*) (Jha et al., 2021). Thus, we conclude that the *RDR3* copies were lost in some gymnosperm lineages, with as-yet unknown roles in earlier lineages and angiosperms.

Sister to *RDR6*, two copies of *RDR1/2*-like genes were found in most non-seed land plants before these genes diverged to RDR1 and RDR2 in gymnosperm and descendant species, resulting in a tree topology of (A,(N-S,((G1,(Mo1,(Ma1,E1))),(G2,(Mo2,(Ma2,E2)))) (Figure 2). Thus, we hypothesize that an ancestral *RDR* gene was duplicated in the early land plant lineages and diverged to give rise to *RDR1* and *RDR2* in seed plants, which became specialized to generate dsRNA molecules from viral or heterochromatic single-strand RNAs (Herr et al., 2005; Garcia-Ruiz et al., 2010). Adjacent to the RDR1/2/6 clade, we observed one RDR in each of the three filamentous green algae (*Chara braunii*, *Interfilum paradoxum*, *Klebsormidium nitens*). Notably, genomes of the filamentous green algae each encode one RDR6 protein, and every land-plant genome encodes one or more RDR6 proteins. Together, these results suggest that RDR1/2/6 diverged before the split between green algae and land plants, resulting in an RDR6 protein tree with a topology of (A,(N-S,(G,(Mo,(Ma, E))))), whereas RDR1 and RDR2 are not found ubiquitously in land plants (but they have diversified in seed plants). The broad presence of RDR6 in land plants is consistent with their highly conserved 21-nt *trans*-acting siRNAs (tasiRNAs) pathway (Xia et al., 2017). RDR6 is also known to play a role in the production of reproductive phasiRNAs; the 21-nt phasiRNAs likely emerged in seed plants (Pokhrel et al., 2021), while the 24-nt phasiRNAs emerged in angiosperms (Xia et al., 2019). These findings suggest that RDR6 became specialized for the tasiRNA pathway prior to the origin of seed plants, and became involved in the reproductive phasiRNA pathways in seed plants.

### The divergence of DCLs and miRNA/siRNA pathways in Viridiplantae

To gain insights into the evolution of DCL proteins and the major sRNA types such as miRNA (processed by DCL1), hc-siRNAs (processed by DCL3), and phasiRNAs (processed by DCL4 or DCL5), we constructed a maximum likelihood tree of 1,036 DCL proteins (including 12 non-plant DCLs). Three distinct clades (DCL1, DCL2/4 and DCL3/5) were observed in Viridiplantae, with DCL3/5 being sister to DCL2 and DCL4, and DCL1 being sister to DCL3/5 and DCL2/DCL4 (i.e., a tree topology of (DCL1,(DCL3/5,(DCL4,DCL2))).

DCL1 was found in the two filamentous green algae and all land plants (Figure 3, Supplemental Data 3) with a species tree topology of (A,(N-S,(G,((Ma,Mo),E)))). Similarly, DCL4 was found in the most ancestral land plant species sampled, two *Marchantia* species) and all descendant lineages (Figure 3) with a species tree topology of (N-S,(G,(Mo,(Ma,E)))). Together, these results indicate an early divergence of these DCL proteins, and suggest the existence of the miRNA pathway in filamentous algae and descendant lineages while the 21-nt tasiRNA/phasiRNA pathways emerged in early land plant species. The *DCL2* copies were only detected in seed plants with an evolutionary pattern of (G,(Mo,(Ma,E))) suggesting that DCL2, and potentially the DCL2- derived 22-nt siRNAs, emerged in seed plants. The most ancient lineages in which we detected *DCL3* copies are the two *Marchantia* species following an evolutionary pattern of (N-S,(G,(Ma,(E,Ma)))). This observation suggests that DCL3 arose in the common ancestor of land plants, implying a functional conservation of the hc-siRNA pathway in early land plant species.

**Figure 3.**
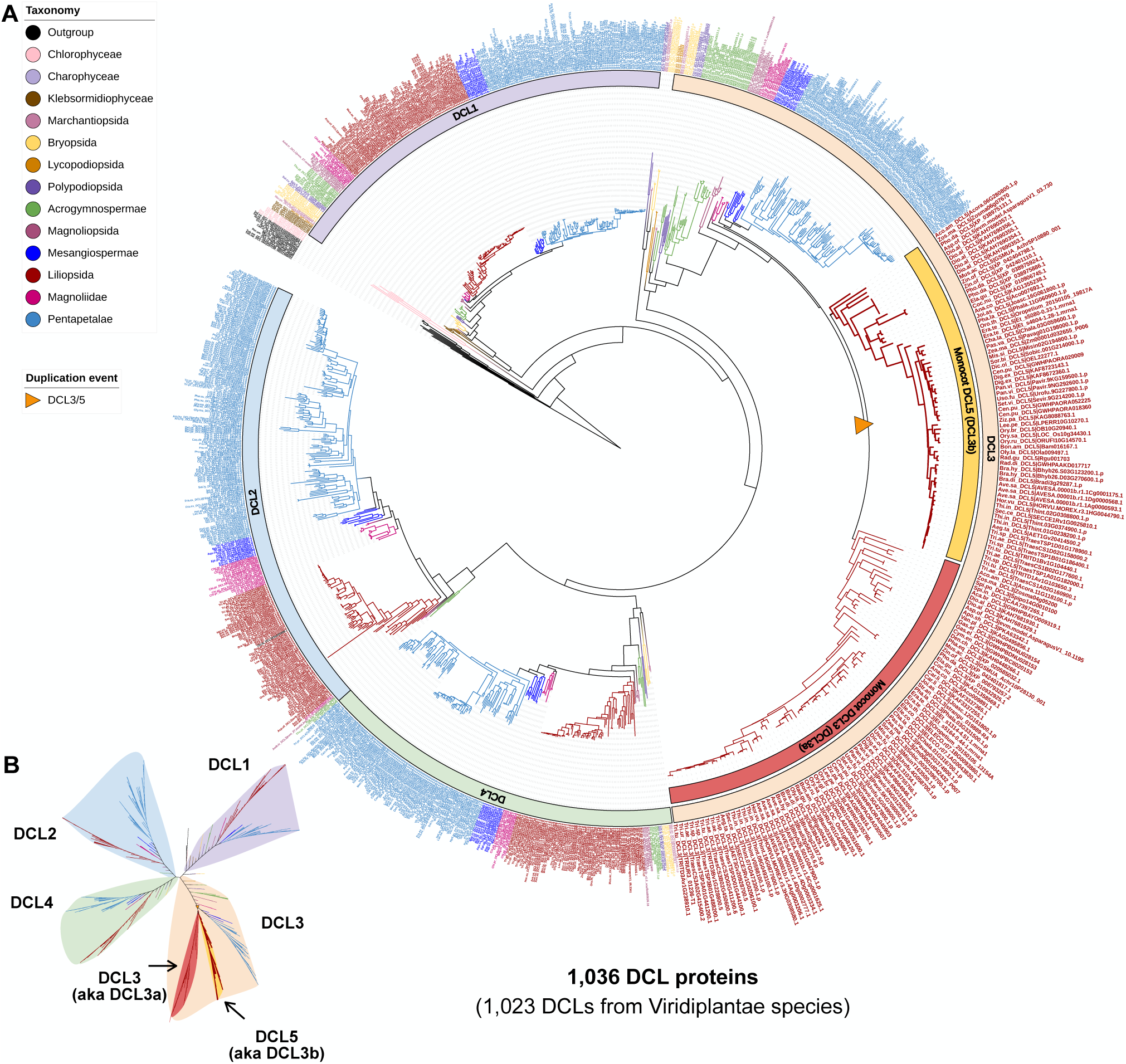
Maximum-likelihood phylogeny of all DCL proteins annotated in the analyzed genomes. **(A)** Root complete phylogeny of all DCL-type proteins in the 207 plant and non-plant species. **(B)** Unrooted view of the major clades of the phylogenetic tree in (A).

DCL5, initially named DCL3b, catalyzes the biogenesis of 24-nt reproductive phasiRNAs (Song et al., 2012; Arikit et al., 2013; Teng et al., 2020; Patel et al., 2021). 24-nt reproductive phasiRNAs were found to peak in abundance at the meiotic stage of anthers in rice and maize (*Zea mays*) (Zhai et al., 2015b; Fei et al., 2016), and more recent work has identified a group of 24-nt phasiRNAs that are highly abundant in pre-meiotic anthers in barley and wheat (Bélanger et al., 2020). The maize loss-of-function *dcl5* mutant fails to produce 24-nt reproductive phasiRNAs but not hc-siRNAs (Teng et al., 2020), demonstrating its specialized function in the reproductive phasiRNA pathway. DCL5 was described as monocot-specific and only present in *Dioscorea* and more recently diverged monocot lineages, although it appeared to have been lost independently in some orders (Patel et al., 2021). Although eudicot genomes do not encode the canonical DCL5 protein, some eudicots also accumulate 24-nt reproductive phasiRNAs, likely generated by DCL3, which is known to catalyze biogenesis of 24-nt siRNAs (Xia et al., 2019; Patel et al., 2021). It was previously hypothesized that DCL5 emerged from DCL3 via whole-genome duplication (WGD) or a *DCL3* gene duplication some time before the diversification of *Dioscorea,* and became specialized for the biogenesis of 24-nt reproductive phasiRNAs by neofunctionalization (Patel et al., 2021). Our current data extends previous findings by detecting a *DCL5* copy in *Asparagus officinalis*, *Phoenix dactylifera*, *Zostera marina*, and *Acorus americanus*, which are monocot species that diverged earlier than *Dioscorea; A. americanus* is also known to be the most ancestral extant lineage of monocots (Duvall et al., 1993). The evolutionary pattern of DCL3 and DCL5 that can be summarized as (N-S,(G,(Ma,(E,(Mo1,Mo2)))))) (Figure 3). Notably, the previous study (Patel et al., 2021) did not detect a *DCL5* copy in the *A. officinalis* genome, whereas we detected one in the current study. This is possibly due to the different and potentially more robust computational approaches used here.

The basal-most angiosperms, including one Amborellales (*Amborella trichopoda*) and three Nymphaeales (*Euryale ferox*, *Nymphaea colorata* and *Nymphaea thermarum*), cluster on a branch that is well supported (bootstrap of 100) and separated from monocots and eudicots. 24-nt reproductive phasiRNAs are found in *Amborella trichopoda* (Xia et al., 2019). However, we found only one *DCL3* copy but no *DCL5* in the *Amborella* genome (Figure 3). This is consistent with a previous study suggesting a dual functionality of DCL3 in the biogenesis of 24-nt phasiRNAs and hc-siRNAs in some extant eudicots (Xia et al., 2019). Notably, the Nymphaeales species have two copies of *DCL3*. Thus, as an alternative hypothesis, *DCL3* was possibly duplicated in basal-most angiosperm and had a dual function to process hc-siRNAs and 24-nt phasiRNAs. Then magnoliids and eudicots lost one *DCL3* copy during the speciation of core angiosperms while the monocots retained the two paralogs. In monocots, *DCL3* copies subfunctionalized resulting in functional divergence of DCL3 and DCL5 to specialize in hc-siRNA and 24-nt phasiRNA pathways, respectively. Future functional and biochemical studies of DCL3 and/or DCL5 across diverse angiosperms (magnoliids; non-commelinid monocots; basal and core eudicots) would be necessary to elucidate the functional divergence of DCL3/5; for example, the *Acorus* DCL3/5 copies could be assessed for functional divergence and expression patterns to determine whether DCL5 properties were distinct at the earliest point in monocots.

### Extensive expansion and divergence of the AGO family in green plants

To characterize the evolution of AGO family proteins in green plants, we constructed a maximum likelihood phylogeny of 2,979 AGO proteins (including 52 non-plant AGOs) (Supplemental Figure 1; Supplemental Data 4; Supplemental Table 2). Consistent with previous reports, we found three major clades, i.e. the AGO4/6/8/9, AGO2/7 and AGO1/5/10/18 clades (Figure 4, 5 and 6; Supplemental Data 4; Supplemental Figure 1).

**Figure 4.**
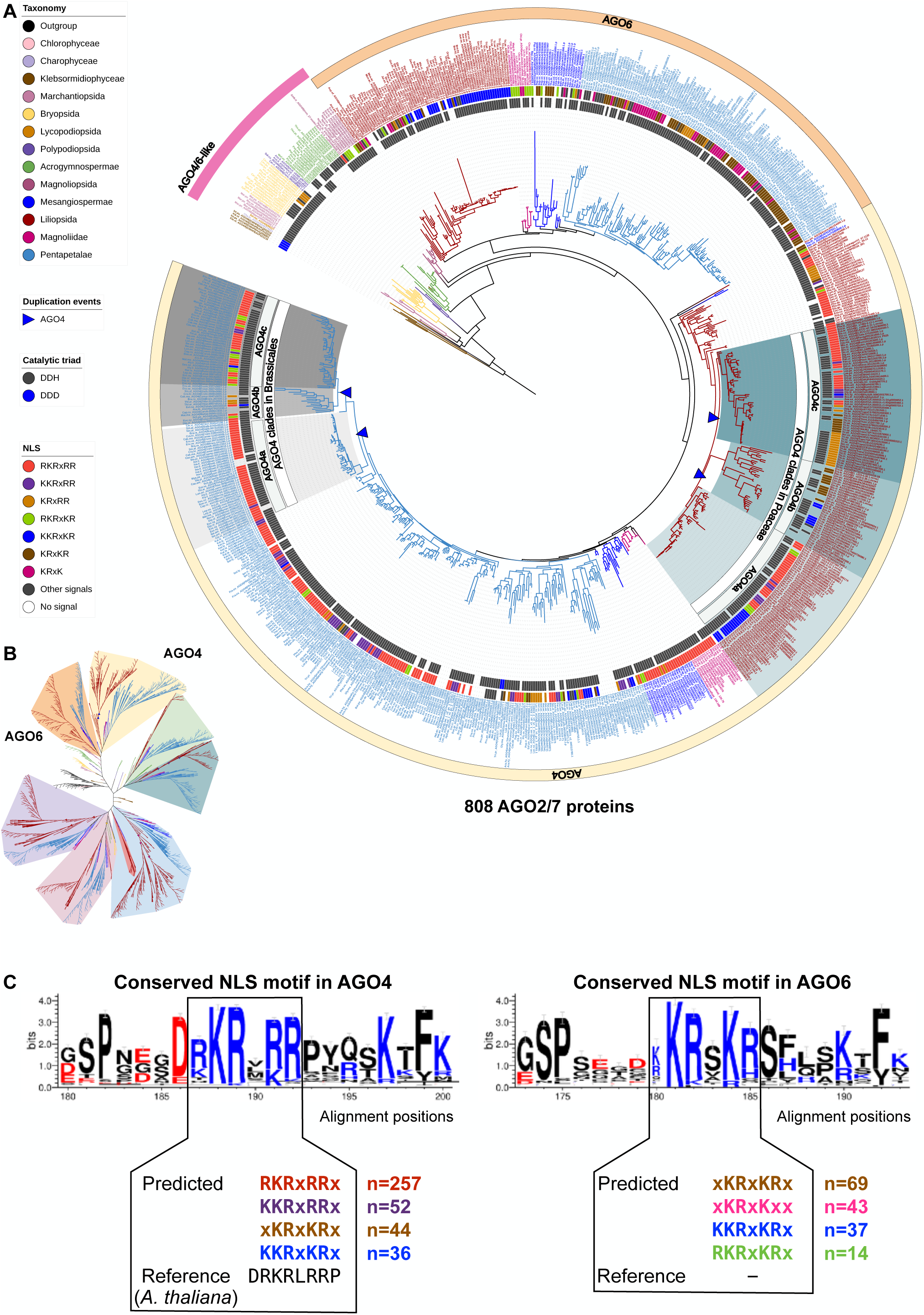
Maximum-likelihood phylogeny of AGO4/6 clade proteins annotated in the analyzed genomes. **(A)** The AGO4/6 clades pruned from the complete AGO phylogeny shown in Supplemental Figure 1. **(B)** Unrooted view of the major clades of the complete AGO tree, with only the AGO4/6 clades highlighted. **(C)** Conserved NLS motifs in the AGO4/6 proteins.

**Figure 5.**
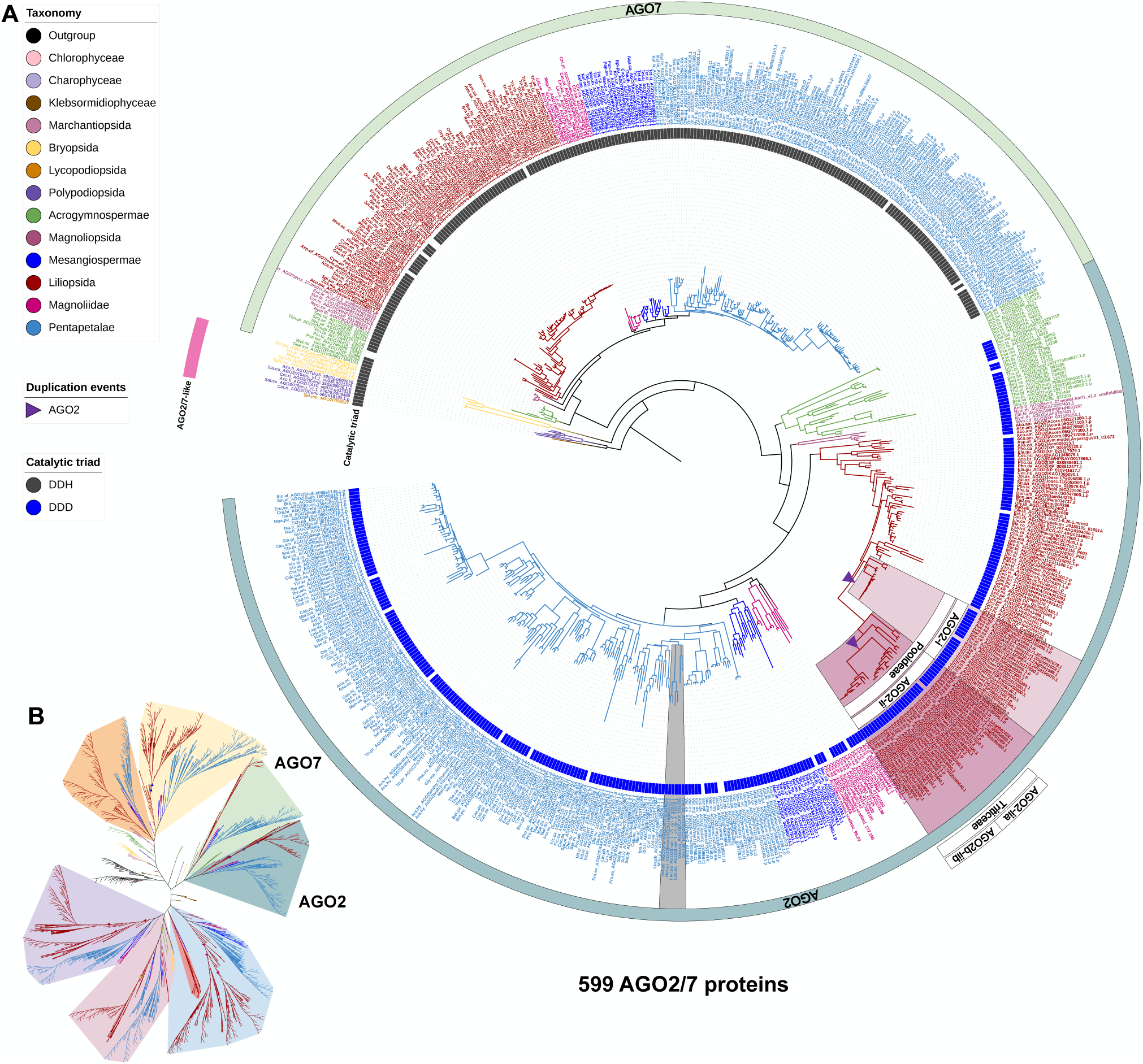
Maximum-likelihood phylogeny of AGO2/7 clade proteins annotated in the analyzed genomes. **(A)** The AGO2/7 clades pruned from the complete AGO phylogeny shown in Supplemental Figure 1. **(B)** Unrooted view of the major clades of the complete AGO tree, with only the AGO2/7 clades highlighted.

The number of *AGO* copies per genome varies among plant lineages, ranging from 1.5 copies on average in green algae to 21.0 copies on average in monocots (Table 2). A recent study described a phylogenetic analysis of 2,958 AGO proteins from 244 plant species (Li et al., 2021a). Although we analyzed 48 fewer plant species, in each of the lineages summarized in Table 2 (except Lycopodiopsida), we identified a larger number of AGO proteins (Li et al., 2021a), suggesting that our work identified a more complete set of AGO proteins. For instance, we found many more AGO proteins encoded in genomes of *Hordeum vulgare*, *Sorghum bicolor*, *Zea mays* and *Triticum aestivum*, with 21, 18, 21, and 75, respectively, compared to 12, 15, 17, and 40 identified by Li et al. (2021b). Such differences are likely due to (i) the source of genome annotation, in our case prioritizing Ensembl Genomes and Phytozome databases, rather than NCBI, (ii) the sensitivity of the method we used to identify orthologous AGO proteins (e.g., the use of OrthoFinder), and (iii) the addition of recently sequenced large monocot genomes (e.g., *Secale cereale*, which encodes 23 AGO proteins).

**Table 2.**
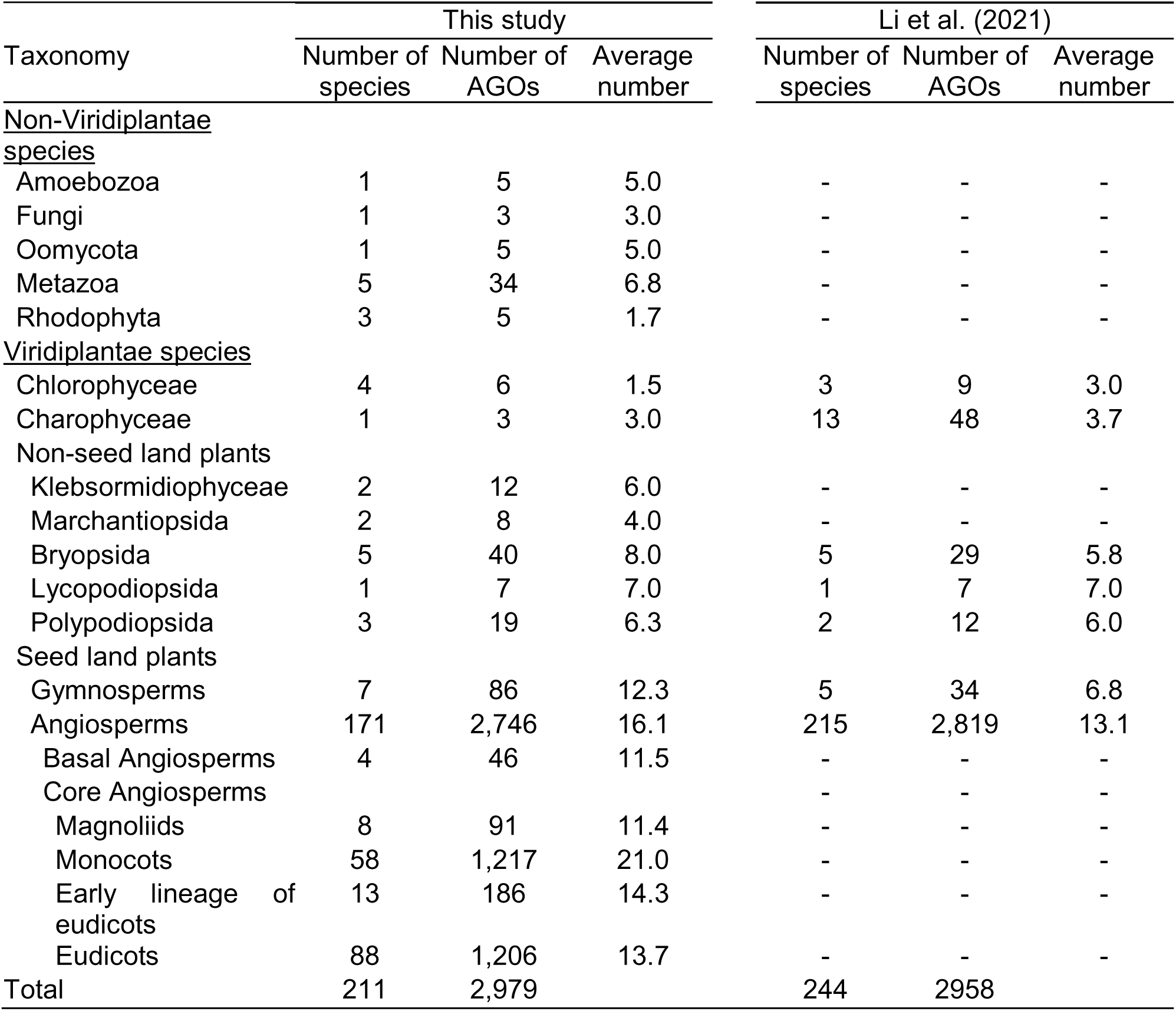
Comparison of the numbers of AGO proteins annotated in this study versus the Li et al. (2021) study.

### Distinct patterns of AGO4 expansion in monocots versus eudicots

Members of the AGO4/6/8/9 clade are best known as effectors of hc-siRNAs (Zhang et al., 2015). Our phylogenetic tree contained, in total, 57 AGO-like proteins in the filamentous green algae, liverwort, lycophyte, moss, fern, gymnosperm and basal-most angiosperm lineages (Supplemental Figure 1), whereas in all flowering plants descendant to Nymphaeales, the AGO family expanded and diverged extensively, resulting in groups of AGO4 and AGO6 (Figure 4; Supplemental Data 4), with a evolutionary pattern of (N-S,(G,((Mo,(Ma,E),(Mo,(Ma,E))))) for (AGO4/6-like,(AGO4/6- like,((AGO6,(AGO6,AGO6),(AGO4,(AGO4,AGO4))))). AGO protein names have been based largely on phylogenetic studies of Arabidopsis, and thus distinct names (AGO4/6/8/9) were assigned to four related AGO proteins found in the same clade. These names were previously applied to rice, maize, and other species (Zhang et al., 2015), which in some cases generates confusing relationships. Thus, we propose that the clade previously known as AGO4/6/8/9 should be renamed as the AGO4 and AGO6 clades, which are two clearly distinguishable clades in flowering plants, whereas AGO8 and AGO9 appear to be paralogs of AGO4 that arose in the common ancestor of Brasicales (Figure 4A).

AGO proteins in Amborellales (*Amborella trichopoda*) and Nymphaeales (*Euryale ferox*, *Nymphaea colorata* and *Nymphaea thermarum*), which are four basal-most angiosperms, forms a clade that is separate from the AGO4 and AGO6 clades that include AGO4/6 proteins of core angiosperms, while eight Magnoliid, two Chloranthales and 11 basal eudicots have AGO proteins in both clades (Figure 4). These results indicate that AGO4 and AGO6 diverged in a common ancestor of core angiosperms. The AGO6 proteins formed a monophyletic clade including all core angiosperms, such that the protein tree mirrors the species tree (Figures 1 and 4), suggesting that AGO6 function is highly conserved within angiosperms. In contrast, the AGO4 proteins display more complex evolutionary patterns (Figure 4A), suggesting that the *AGO4* genes have undergone extensive duplication and possibly neofunctionalization/subfunctionalization. The distinct evolutionary patterns of AGO4 versus AGO6 are in line with the diverged expression patterns and functions of these two clades of proteins, although they have been shown to be functionally redundant in several studies (Havecker et al., 2010; Duan et al., 2015).

Within monocots, AGO4 exhibits distinct evolutionary patterns in Poaceae and non-Poaceae species. We observed lineage-specific *AGO4* duplication events in several species within the Zosteraceae (species *Zostera marina*), Asparagaceae (*Asparagus officinalis*), Orchidaceae (*Dendrobium catenatum*, *Phalaenopsis equestris*) families, resulting in two *AGO4* copies per genome. a Bromeliaceae (*Ananas comosus*) and Cyperaceae (*Carex littledalei*), two Poales species, each has two *AGO4* copies clustered species-by-species, suggesting independent gene duplication events (Figure 4A). In contrast, three *AGO4* copies were detected in nearly every Poaceae species, forming three monophyletic groups (Figure 4), which may be the result of an *AGO4* gene duplication event in the common ancestor of the Poaceae family. We hypothesize that the different number of gene duplication events observed in the AGO4 and AGO6 clades is likely to be associated with the divergence of the sRNA pathways in which they are involved. For example, it has been suggested that AGO4 and AGO6 are specifically involved in the RNA-directed DNA methylation pathway, but have no clear developmental roles in Arabidopsis, whereas at least one member (in maize) of the AGO4c clade of Poaceae is known to play an important role in regulating reproductive development (Singh et al., 2011). Together, our phylogenetic data, and the published functional data, suggest multiple gene duplication events and neofunctionalization specific to the *AGO4* genes in monocots.

The evolution of AGO4 in eudicots is even more complex as the AGO4 group expanded several times independently. For instance, in the Brassicales, we observed three AGO4 subgroups (named Brassicales AGO4a, AGO4b and AGO4c) that form a single larger clade distinct from other species in the Malvids group (Figure 4A). The *AGO4* copies in the Brassicales-AGO4b clade (represented by the Arabidopsis *AGO8*) appears to be duplicated from the genes in the Brassicales-AGO4a clade (represented by the Arabidopsis *AGO4*), while another gene duplication event may have generated the *AGO4* copies in the *AGO4c* group (represented by the Arabidopsis *AGO9*). All Brassicales species have one or more *AGO4a* and *AGO4c* copies, while several Brassicales species are absent from the AGO4b clade. The Arabidopsis *AGO8* gene is found in this clade and has been suggested as a pseudogene in Arabidopsis (Mallory and Vaucheret, 2010).

Arabidopsis AGO4 is known to function in the RNA-directed methylation pathway, with both cleavage-dependent and independent roles (Qi et al., 2006; Wang and Axtell, 2017). To determine if mediating RNA cleavage is a conserved role of AGO4/6 proteins, we identified the catalytic triads of the PIWI domain of all proteins in this clade. Most of the proteins in land plants have a conserved Asp-Asp-His [DDH] triad, except a few that have an Asp-Asp-Asp [DDD] triad (Figure 4A). Notably, all the AGO4/6-like proteins in the filamentous green algae have a DDD triad, which is commonly observed in AGO2 proteins of seed plants (see below). These data suggest that a one-amino acid change in the catalytic triad occurred in the common ancestor of the green alga *Chara braunii* and land plants, and that most, if not all, of the AGO4/6 proteins in green plants have the potential to mediate RNA cleavage. We also analyzed the AGO4/6 proteins for NLS sequences. We detected an NLS nearly ubiquitously in the AGO4/6 proteins of green plants (Figures 4A). A close inspection of the NLS sequences in AGO4 and AGO6 demonstrated that the sequences are similar between the two clades of proteins, and are all localized near the N terminus (Figure 4C). Together, these results support conserved roles of many plant AGO4/6 proteins in the RNA-directed DNA methylation pathway (Borges and Martienssen, 2015).

### AGO2/7 emerged in mosses and diverged in seed plants

Members of the AGO2/7 clade are known to function in bacterial/viral defense (AGO2) or tasiRNA biogenesis (AGO7) (Harvey et al., 2011; Zhang et al., 2015). The AGO2/7 clade of our AGO tree shows a group of 12 AGO2/7-like proteins present in moss, lycophyte and fern species that then diverged and expanded in seed plants, resulting in two groups: AGO2 and AGO7 (Figure 5; Supplemental Table 2). We did not identify AGO2/7-like genes in the genomes of any of the filamentous algae or liverwort species, indicating that the AGO2/7 clade emerged in non-vascular plants and then diversified in seed plants. We infer that AGO7 emerged in seed plants, as we identified one AGO7 ortholog in most gymnosperms, and a monophyletic AGO7 group spanning all flowering plants. AGO7 catalyzes the miR390-directed cleavage of *TAS3* transcripts, yielding 21- nt tasiRNAs regulating Auxin response factor (ARF) family transcription factors in angiosperms (Montgomery et al., 2008). Previous studies reported a miR390-*TAS3*- ARF regulatory module acting in the bryophyte *Physcomitrella patens*, and in some later diverged lineages, in the absence of AGO7 protein (Axtell et al., 2006; Axtell et al., 2007; Arif et al., 2012; Cho et al., 2012; Liu et al., 2020). Thus, we hypothesize that the moss AGO2/7-like protein might have already evolved a function that is analogous to canonical AGO7 proteins and is involved in the biogenesis of tasiARFs in bryophytes, while the miR390-triggered AGO7-catalyzed *TAS3*-ARF regulatory module was completed and refined in the course of the evolution of seed plants.

AGO2 likely diverged from AGO7 in the common ancestor of seed plants, as genomes of the gymnosperm, basal angiosperm and core angiosperm lineages encode AGO2, while moss, lycophyte and fern genomes encode an AGO2/7-like protein. In monocots, subgroups of plant species appear to have lost their *AGO2* gene copy as genomes of Alismatales (two Araceae and one Zosteraceae species), Asparagales (six Orchidaceae species), Dioscoreales (one Dioscoreaceae species), two Zingiberales (one Musaceae and one Zingiberaceae) and Poales (one Cyperaceae) species do not encode an AGO2 protein (Figure 5). The *AGO2* gene was apparently duplicated in a common ancestor of the Poaceae species, resulting in two *AGO2* clades in Poaceae species (Figure 5). An analysis of catalytic triads in AGO2 and AGO7 proteins showed that all the AGO7 proteins in seed plants have the catalytic triad sequence DDH, whereas the AGO2 proteins in seed plants all have a DDD triad (Figure 5). Since the AGO2/7-like proteins in bryophytes, lycophytes, and ferns also have the DDH triad, an H to D change likely occurred in the common ancestor of seed plants, perhaps concurrently with the functional divergence between AGO2 and AGO7.

AGO2 proteins in eudicots did not cluster in monophyletic groups, suggesting that they did not originate in a common ancestor of eudicots. As an example, we observed a complex and ambiguous evolution of *AGO2* genes in the genomes of Asterids, with two AGO2 proteins in each of two Asterales (*Helianthus annuus* and *Lactuca sativa*) and one Apiales (*Daucus carota*). All these AGO2 proteins are on the same branch and grouped by genus. Together, the evolution of *AGO2* genes in the Poaceae and eudicots exhibit complex and species/genus-specific patterns, which might be consequences of co-evolution with various bacterial/viral pathogens.

### Extensive expansion of the AGO1/5/10/18 clade in monocots

AGO1 and AGO10 proteins are canonical effectors of miRNAs (Zhang et al., 2015). Our AGO tree shows that liverworts, mosses, lycophytes, ferns and gymnosperms cluster with AGO10 but not AGO1 (Figure 5; Supplemental Figure 1; Supplemental Data 4), and that AGO1 diverged from AGO10 in early lineages of flowering plants, as AGO1 was observed in basal-most lineages and descendants. A recent phylogenetic analysis of plant AGO proteins detected a similar feature (Li et al., 2021a). These observations suggest that non-seed species have a single AGO10 protein that acts as the predominant effector of miRNAs. In Arabidopsis, AGO10 regulates shoot apical meristem development by sequestering miR165/6 and promoting their degradation, preventing them from being loaded onto AGO1. AGO1 regulates development and stress responses, and AGO18 plays a role in antiviral response (Zhang et al., 2015).

Thus, our results suggest that AGO10 emerged first and then diversified to AGO1/5/18 in flowering plants to play distinct roles. The evolution of AGO10 has otherwise been well described, including a monocot-specific duplication of *AGO10b* (Figure 6) (Li et al., 2021a). Thus, we focus on the evolution of AGO1, mainly in monocots. Our phylogenetic tree indicates that a duplication of *AGO1* occurred specifically in the monocot lineage soon after the monocot/eudicot split, resulting in two monophyletic groups, which we refer to as AGO1-i and AGO1-ii (Figure 6). Then, another duplication event of each group occurred in a common ancestor of the Poaceae to generate two sub-groups, resulting in four groups, called Poaceae-AGO1-i to AGO1-iv (Figure 6). *AGO* gene duplication or loss events seem to have occurred in a lineage-specific manner. First, the group Poaceae-AGO1-i represents most Poaceae species, except some PACMAD species like maize and sorghum. Second, all PACMAD species are absent from the AGO1-iii group. Notably, the *AGO1-iii* copies are further duplicated in BOP species generating a unique clade of proteins, represented by AGO17 in rice (*Oryza sativa*), but many BOP species lost this AGO after the duplication event. Furthermore, AGO17 has been suggested to be rice specific (Pachamuthu et al., 2021), but our analysis identifies an AGO17 protein in at least two other *Oryza* species (*O. glaberrima* and *O. rufipogon*) and non-*Oryza* species of Poaceae (*Eragrostis tef, Miscanthus sinensis, Dichanthelium oligosanthes, Cenchrus purpureus*, and a few *Triticum* species). These results refine the origin and evolution of the AGO17 protein, whose evolutionary pattern has been controversial – sometimes found outside the AGO1 clade (Zhang et al., 2015; Pachamuthu et al., 2021), and sometimes within the AGO1 clade (Bélanger et al., 2020; Li et al., 2021a). Functional studies of AGO17 have demonstrated its roles in regulating rice development and loading miR397 (Zhong et al., 2020; Pachamuthu et al., 2021). In line with these studies, our data suggest a potential redundancy between AGO17 and AGO1 (Zhong et al., 2020). Finally, duplicated *AGO1- iv* copies are observed in all PACMAD species and Triticeae species but absent from Brachypodieae genomes, suggesting a loss of these copies specifically in the Brachypodieae lineage. As the canonical effectors of miRNAs, the expansion of AGO1 clade in monocots might be associated with the evolution of specialized sRNA pathways involving miRNAs, e.g. the reproductive phasiRNA pathways that utilize miRNAs as triggers of phasiRNAs.

**Figure 6.**
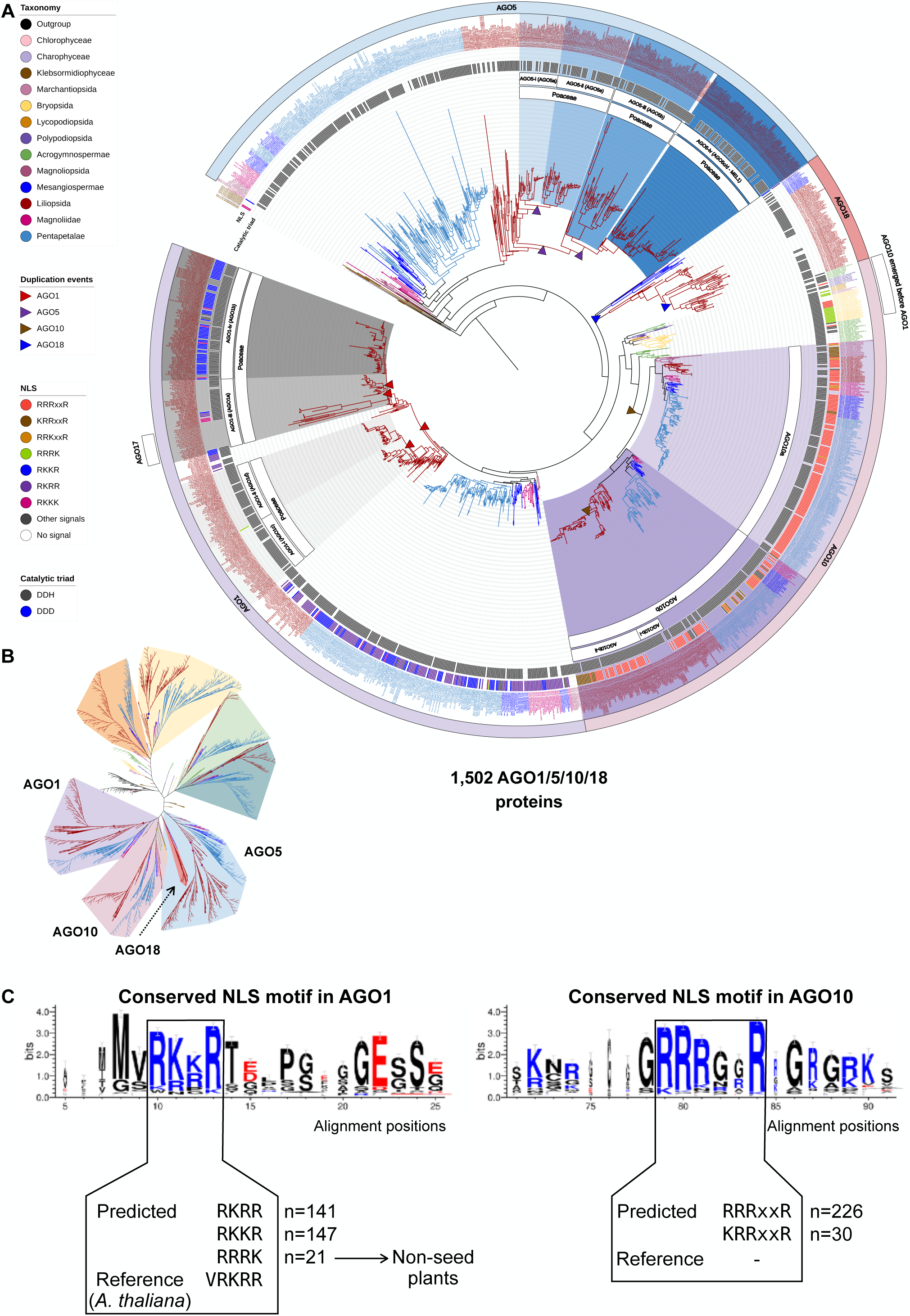
Maximum-likelihood phylogeny of AGO1/5/10/18 clade proteins annotated in the analyzed genomes. **(A)** The AGO1/5/10/18 clades pruned from the complete AGO phylogeny shown in Supplemental Figure 1. **(B)** Unrooted view of the major clades of the complete AGO tree, with only the AGO1/5/10/18 clades highlighted. **(C)** Conserved NLS motifs in the AGO1/10 proteins.

AGO18 has been described as a Poaceae-specific clade (Zhang et al., 2015; Li et al., 2021a), with a known role in virus resistance (Wu et al., 2015), and has been proposed as a candidate effector(s) of reproductive phasiRNAs in anthers (Zhai et al., 2015b; Fei et al., 2016; Bélanger et al., 2020; Das et al., 2020). Within Poales, we found an AGO18 protein in *Carex littledalei* (Cyperaceae) and *Joinvillea ascendens* (Joinvilleaceae) but not in *Ananas comosus* (Bromeliaceae). In addition, in *Amborella trichopoda,* a basal-most angiosperm, and all basal eudicots, we detected an AGO protein that clusters with the Poales AGO18 proteins. These data indicate that AGO18 emerged in the common ancestor of angiosperms; this contrasts with previous studies suggesting that AGO18 is grass specific (Zhang et al., 2015; Li et al., 2021a). However, all the non-commelinids monocots, magnoliids, and core eudicots have lost AGO18, suggesting that the role of AGO18 could be compensated for by other AGO proteins in these lineages. Additionally, our data demonstrate that the AGO18 clade clusters with AGO1/10, contrasting with the previous studies suggesting the AGO18 clade being sister to the AGO1/5/10 clade (Zhang et al., 2015). Therefore, our analyses refine the origin and evolution of AGO18. A few studies have demonstrated crucial roles of AGO18 in reproductive development and antiviral defense (Wu et al., 2015; Sun et al., 2018; Sun et al., 2019), but future work would be necessary to understand why AGO18 is retained (and duplicated) in grasses yet was apparently lost in the majority of other angiosperm lineages.

The AGO5 proteins are known to be angiosperm specific (Li et al., 2021a a). The number of AGO5 proteins greatly expanded in Poaceae species, and the evolutionary pattern of AGO5 is more complicated than that of AGO1, AGO2 or AGO4. Poales AGO5 proteins diverged from *Musa* and other monocot lineages before expanding in Poaceae species, resulting in three major groups, which we refer to as Poaceae-AGO5- i, ii, and iii (Figure 6; Supplemental Figure 1; Supplemental Data 4). Some species have additional *AGO5* copies in the Poaceae-AGO5-i clade. AGO5-i diverged twice in the common ancestor of Poaceae, resulting in three branches where (i) one branch is represented by Oryzoideae species only, (ii) the second branch is represented by Pooideae species only and (iii) the third branch includes all Poaceae species (Figure 6). This pattern of AGO5-i divergence demonstrates gene duplication events that result in a gain of an *AGO5* copy in Oryzoideae and Pooideae lineages. Similarly, we observed two *AGO5* copies in Oryzoideae species of Poaceae-AGO5-ii while only one in all other Poaceae, suggesting another duplication of *AGO5* in the Oryzoideae group (Figure 6). Interestingly, one of the AGO5-iii group genes encodes the rice AGO5c protein, also called MEIOSIS ARRESTED AT LEPTOTENE 1 (MEL1), and is known as a binding partner of 21-nt reproductive phasiRNAs and essential for the progression of meiosis in anthers (Nonomura et al., 2007; Komiya et al., 2014). Similarly, in maize, *AGO5c* has recently been identified as the causal gene of the classic mutant *male sterile 28* (*ms28*) and is essential for anther development and male fertility (Li et al., 2021b). Separate work demonstrated that AGO5b and AGO5c in maize, named as MALE-ASSOCIATED ARGONAUTE-1 (MAGO1) and -2 (MAGO2), respectively, play redundant roles in loading 21-nt phasiRNAs in pre-meiotic anthers to regulate retrotransposons (Lee et al., 2021). These data suggest a conserved role of AGO5 proteins in the premeiotic, 21-nt phasiRNA pathway. Notably, MAGO1 is a member of the AGO5-iii clade while MAGO2 is in the AGO5-ii clade, suggesting functional redundancy between the two clades.

An analysis of the catalytic triads in the AGO1/5/10/18 proteins showed that except for the algal clade (*Klebsormidium nitens, Interfilum paradoxum, and Chara braunii*), a DDH triad is found nearly all clades of these AGO proteins (Figure 6), suggesting that this group of AGO proteins in land plants evolved the ability to mediate RNA cleavage after the divergence between algaes and land plants, and since that divergence, the catalytic triad has been highly conserved. One clade of AGO18 proteins, encoded by genes that were duplicated specifically in a few monocot species, does not have a conserved catalytic triad (Figure 6), suggesting that these AGO18 paralogs evolved new functions other than mediating transcript cleavage. An analysis of the NLS in the AGO1/5/10/18 clade proteins showed that none of the AGO5 or AGO18 clade proteins has an NLS. Among the AGO1 proteins, those in the AGO1-iv clade all have an NLS, but those in the AGO1-i/ii/iii clade do not, indicating that the AGO1-iv proteins are more likely to function in the nucleus, and suggesting a functional divergence among the four clades of AGO1 proteins derived from the Poaceae-specific gene duplication events. Similarly, the AGO10b-i proteins, which diverged from the AGO10b-ii proteins after a monocot-specific *AGO10* duplication event, all lack an NLS (Figure 6A), suggesting functional divergence of AGO10b-i versus AGO10b-ii proteins. The NLS sequences are distinct between AGO1 and AGO10, but they are highly conserved in each of the two clades (Figure 6C).

Together, our analyses of the AGO1/5/10/18 clade suggest that (i) green algae genomes do not encode any AGO protein in the AGO1/5/10/18 clade, known as the effector of miRNAs, (ii) *AGO10* emerged in moss and expanded in flowering plants, yielding more *AGO10* copies, and the *AGO1* copies, (iii) only angiosperm genomes encode AGO5, and (iv) AGO18 emerged in the basal-most angiosperm but was lost in most lineages except basal eudicots and Poales. Moreover, our data demonstrate that *AGO1/5/18* gene copies increased in a common ancestor of Poaceae and then was followed by lineage-specific gene gain or loss events. Since AGO1, AGO5, and AGO18 are the only three clades that have been shown to be involved in reproductive phasiRNA pathways, their expansion in the grass lineage may have been driven by a need to form RNA-Induced Silencing Complexes (RISCs) with miR2118 or miR2275 (e.g., AGO1 paralogs), to trigger biogenesis of either 21- or 24-nt reproductive phasiRNAs, or to load the phasiRNAs (e.g., AGO1, AGO5 and AGO18) and act on their targets.

### Evolution of accessory proteins of sRNA pathways

To further understand the evolution of protein complexes involved in sRNA biogenesis and modification, we performed phylogenetic analyses of the DRB and SE (putative protein partners of DCLs), and SGS3 (a putative partner of RDRs), and HEN1 proteins. Our analyses yielded phylogenetic trees for 455 SGS3 (Supplemental Figure 2; Supplemental Data 5), 2,000 DRB (Supplemental Figure 3; Supplemental Data 6), 470 SE (Supplemental Figure 4; Supplemental Data 7), and 224 HEN1 (Supplemental Figure 5; Supplemental Data 8).

Consistent with the role of SGS3 in the plant-specific tasiRNA pathway, no SGS3 proteins were found in any outgroup species (Amoebozoa, Fungi, Metazoa or Oomycota), nor in the Rhodophyta and Chlorophyta species, whereas three filamentous green algae genomes encode SGS3 suggesting that SGS3 emerged in the Viridiplantae genome before the speciation of land plants (Supplemental Figure 2). Interestingly, the genomes of some outgroup species encode an RDR protein (one Amoebozoa, one Fungi, one Metazoa and one Oomycota) suggesting that RDR emerged before SGS3. Although most land plants have two *SGS3* copies per diploid genome, all proteins evolved from the same branch, such that the protein tree mimicked that of the species tree, suggesting that no subfunctionalization or neofunctionalization evolutionary processes occurred in SGS3 paralogs before speciation. The ubiquitous presence of SGS3 in land plants suggests a lack of substrate specificity of SGS3, which in turn suggests that SGS3 proteins do not specialize in certain sRNA biogenesis pathways, but rather are a protein partner to specific RDR proteins, such as RDR2 or RDR6 that are specific to each siRNA biogenesis pathway.

We observed five distinct DRB clades. Compared to the other families that we analyzed, some branches are quite long, in particular, the lengths of the branches separating DRB3 from other DRB proteins and separating monocots and eudicots in DRB3, and one group of duplicated DRB4 proteins in eudicots (Supplemental Figure 3; Supplemental Data 6; Supplemental Table 1). Only one DRB protein has been detected in the outgroup species (Amoebozoa; *Dictyostelium discoideum*), none in rhodophyta nor chlorophyta species, whereas DRB proteins were found in all three filamentous green algae, indicating that Viridiplantae genomes encoded DRB proteins before the emergence of land plants. Rooted and reconciled with the species tree, we observed five DRB clades (DRB1-DRB5) in Viridiplantae, with DRB3 being sister to DRB4 and DRB2, and DRB1 sister to DRB3 and DRB4/DRB2 while DRB5 is adjacent to DRB1 (i.e., a tree topology of DRB5,(DRB1,(DRB3,(DRB4,DRB2)))). The evolution of DRB1, a canonical cofactor of DCL1, is monophyletic as it is present in liverworts and descendent lineages (Supplemental Figure 3). Because DCL1 is present in filamentous green algae, the emergence of DRB1 in land plant suggests that DCL1 might be capable of producing miRNAs in the absence of DRB1 in algae, and then DRB1 emerged in land plants to function as a cofactor of DCL1. The miR390-*TAS3*-ARF regulatory module, which involves 21-nt phasiRNA production, requires DCL4 and, presumably, DRB4 protein as a cofactor (Adenot et al., 2006); however, DRB4 has been found only in flowering plants, while tasiRNAs emerged in mosses (Arazi et al., 2005; Axtell et al., 2006; Talmor Neiman et al., 2006; Axtell et al., 2007). Interestingly, we observed a seemingly similar evolutionary timelapse in the emergence of DCL1/DRB1 and DCL4/DRB4 proteins, which suggests that core proteins emerged before their cofactors and thus the emergence of DRB proteins contributed to an evolutionary sophistication of sRNA pathways that may enhance their biosynthetic efficiency or increase the diversity of sRNA products. Besides DRB1, only the DRB2 clade was observed in all plant lineages. Only the DRB2 clade underwent duplication in angiosperms, yielding DRB2b and DRB2c-type proteins. This is perhaps driven by a requirement for distinct DCL/DRB complexes in biogenesis of diverse sRNAs in angiosperms.

Present in most outgroup species, including Rhodophyta, our phylogenetic analysis shows that SE protein is encoded in the genome of Chlorophyta and land plants (Supplemental Figure 4; Supplemental Data 7; Supplemental Table 1). The Arabidopsis genome encodes only one SE protein, known as a partner of DCL1 in the biogenesis of miRNA (Yang et al., 2006b). Recent work has demonstrated a dual role of SE, in miRNA processing and in RNA decay, as part of the nuclear exosomal complex (Bajczyk et al., 2020); thus, it may be well conserved between outgroup and Viridiplantae species because it has essential functions in miRNA and other sRNA biosynthetic activities. Interestingly, the SE protein family diversified in monocots before the split of Poaceae and other Poales species, resulting in three Poaceae groups, which we named SE-i, SE-ii and SE-iii (Supplemental Figure 4). Since these SE proteins diverged in a common ancestor of Poaceae, they may have evolved separately and undergone subfunctionalization or neofunctionalization in Poaceae.

First identified in plants, HEN1 orthologs have also been discovered in bacteria, fungi, and animals, with known roles in catalyzing 2′-O-methylation of sRNAs (Tkaczuk et al., 2006; Saito et al., 2007; Ji and Chen, 2012). We detected at least one copy of *HEN1* in all land plant species (Supplemental Figure 5; Supplemental Data 8; Supplemental Table 2). *HEN1* has remained a single-copy gene in most species including the monocots, but shows modest expansion in some eudicot lineages. For example, we detected two *HEN1* copies in all Brassicaceae species but not in *Cleome violacea* (Supplemental Figure 5). Thus, the function of HEN1 is likely conserved in land plants, consistent with its universal roles in mediating methylation of miRNAs and siRNAs.

## CONCLUDING REMARKS

Compared to the rapidly growing number of sequenced plant genomes, comprehensive evolutionary studies of proteins associated with sRNA biogenesis and function in plants remain limited. Two recent publications that addressed the emergence of some of the proteins that we have analyzed in our study across ancestral lineages of Embryophyta investigated a total of 36 (You et al., 2017) or 34 (Wang et al., 2021b) species. Those studies uncovered the origin of AGO, DCL, HEN1, RDR, SE and other sRNA biogenesis proteins; however, both studies investigated a limited number of gymnosperm (1) and angiosperm (3) species. Other phylogenetic studies focused on angiosperms and analyzed the evolution of some of the proteins we examine in this work (mainly AGO, DCL and RDR proteins). However, these studies were performed on a small number of closely-related species (Xia et al., 2019; Bélanger et al., 2020; Pokhrel et al., 2020; Patel et al., 2021) and thus a complete evolutionary history could not be elucidated. The evolution of the DCL (DCL3/DCL5 in particular) and AGO (AGO4/6 and AGO1/5/10/18, in particular) proteins was not thoroughly analyzed. Our analyses provide comprehensive annotations and refine the evolutionary history for the AGO/DCL/RDR families and several accessory protein families of plant sRNA pathways (Figure 7), and shed light on the evolution of the various sRNA pathways that involve these proteins. Our results also suggest that the conclusions of functional studies in model species such as Arabidopsis may not be easily translated to other groups of species, as reflected by the existence of lineage-specific proteins such as DCL5 and AGO18. The number of plant genome assemblies is increasing rapidly, and up-to-date bioinformatic tools enable evolutionary studies of tens to hundreds of species, from unicellular organisms such algae to polyploid and large genomes like wheat. Therefore, future studies may continue to refine our understanding of the evolution and function of proteins involved in plant sRNA pathways, including proteins that function in sRNA turnover or with lesser or partial roles in biogenesis.

**Figure 7.**
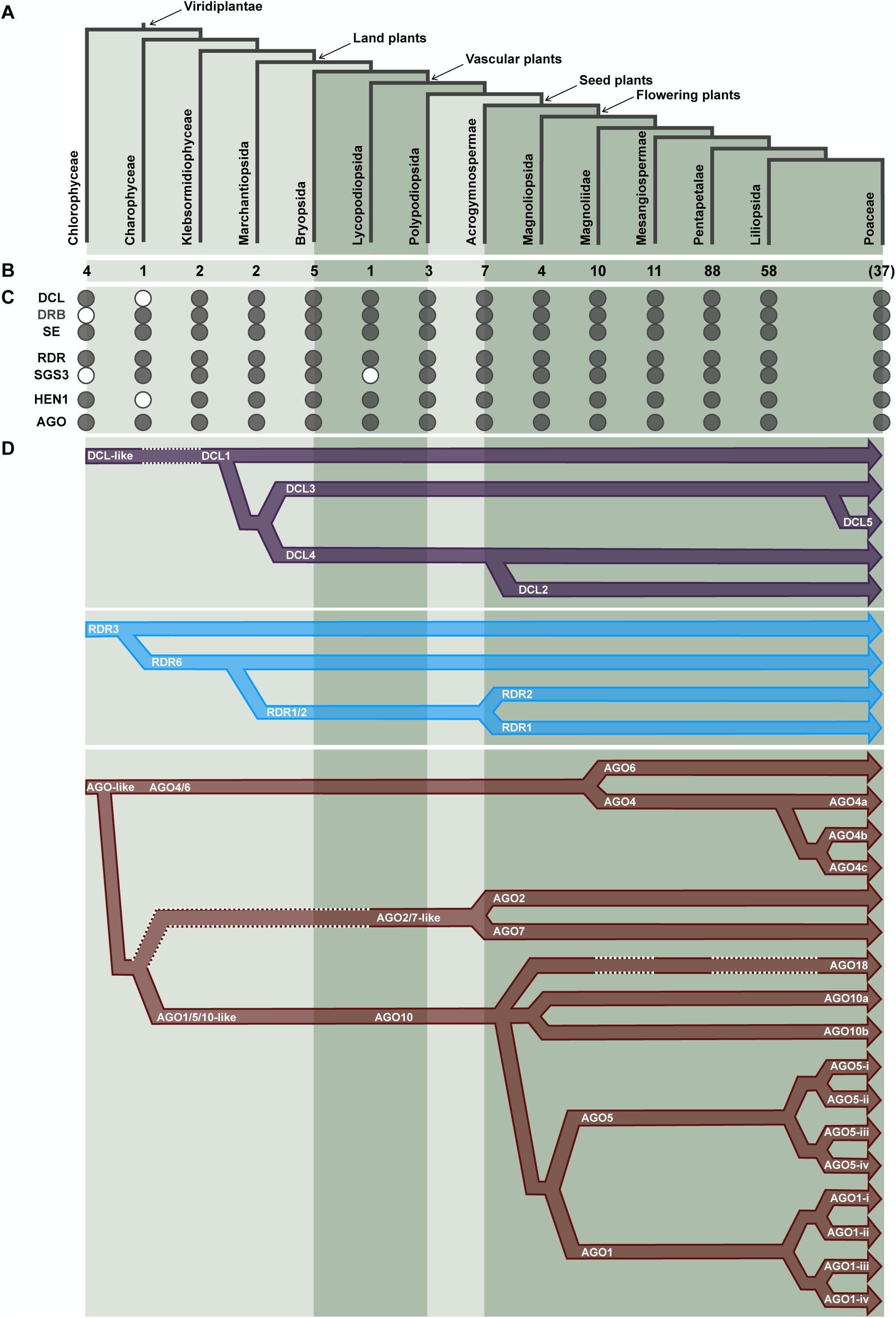
Emergence and diversification of sRNA pathway proteins in plants. **(A)** The major clades of Viridiplantae examined in this study. **(B)** Number of species sampled from each clade. **(C)** Presence or absence of sRNA pathway proteins. **(D)** Refined revolutionary history of DCL/RDR/AGO proteins. Dashed white lines indicate the absence of DCL/RDR/AGO proteins in specific lineages.

## MATERIALS AND METHODS

### Sources of proteome sequences

Proteome sequence files were downloaded from Bamboo genome database (http://www.genobank.org/bamboo), Ensembl Genomes (https://ensemblgenomes.org/), FernBase (https://fernbase.org/), Genome Database for Vaccinium (https://www.vaccinium.org), Genome Warehouse (https://ngdc.cncb.ac.cn/gwh/), GrainGenes (https://wheat.pw.usda.gov/GG3/), Hardwood genomics project (https://www.hardwoodgenomics.org/), Magnoliadb (http://www.magnoliadb.com:7777), NCBI, Phytozome (https://phytozome-next.jgi.doe.gov/), and TreeGenes (https://treegenesdb.org/). A detailed list of all analyzed species is provided in Supplemental Table 1; the taxonomic classification was based on NCBI Taxonomy Database (https://www.ncbi.nlm.nih.gov/taxonomy) and Britanica Encyclopedia (https://www.britannica.com/). Species names were shortened to five-letter codes and combined with protein IDs to label individual proteins in protein trees.

### Ortholog identification and phylogenetic tree construction

OrthoFinder v2.5.4 (Emms and Kelly, 2015) was used to identify orthologous proteins among 207 species using the following parameters: -S diamond -M msa -A mafft -os. Known protein sequences of each family in *Arabidopsis thaliana*, *Chlamydomonas reinhardtii*, *Hordeum vulgare*, *Oryza sativa*, *Physcomitrella patens*, *Glycine max*, *Solanum lycopersicum*, *Triticum aestivum*, *Volvox carteri* and *Zea mays* (reported by (Zhang et al., 2015; You et al., 2017; Bélanger et al., 2020; Baldrich et al., 2022) or available on UniProt database) were used to filter orthologous groups for each protein family. Protein sequences were filtered for the presence of functional domains (Figure 1B) by searching on Pfam r35.0 (Mistry et al., 2020) using hmmscan from HMMER v3.3.2 (http://hmmer.org) with the following parameters: -E 0.00001, and then manually checked/curated using CDvist (Adebali et al., 2015). Unaligned homologous sequences were inspected to identify and remove stretches of non-homologous adjacent characters using PREQUAL v1.02 (Whelan et al., 2018) with default parameters. Multiple sequence alignments (MSAs) were performed using MAFFT v7.505 (Katoh and Standley, 2013; Katoh et al., 2017) with the following parameters: --auto --anysymbol -- dash --originalseqonly. Protein alignments were trimmed with trimAL v1.4.1 (Capella-Gutiérrez et al., 2009) using the following parameters: -gt 0.9 -cons 60 -w 3. Trimmed alignments were used to infer maximum-likelihood phylogenetic trees using IQ-TREE v2.2.0.3 (Minh et al., 2013; Nguyen et al., 2015; Kalyaanamoorthy et al., 2017) with the following parameters: -B 1000 --mset JTT -T AUTO.

### Reconciliation of protein trees with a species tree

To infer a species tree, orthologous groups were identified from the 207 species using OrthoFinder. Multiple sequence alignments (MSAs) were performed using MAFFT v7.505 (Katoh and Standley, 2013; Katoh et al., 2017; Rozewicki et al., 2019) with the following parameters: --auto --anysymbol. Protein alignments were trimmed with trimAL v1.4.1 (Capella-Gutiérrez et al., 2009) using the following parameters: -gt 0.9 -st 0.001 - cons 30 -w 3. Protein trees were inferred using fasttree v2.1.11 (Price et al., 2010) with following parameters: -cat 20 -gamma -spr 4 -sprlength 50 -mlacc 3. A total of 234 protein trees representing 207 species were used to infer the species tree with STAG (Emms and Kelly, 2018). The species tree was rooted using STRIDE (Emms and Kelly, 2017).

Maximum likelihood protein family trees were inferred from their aligned sequences, the mapping between genes and species, and the rooted species tree using GeneRax v2.0.4 (Morel et al., 2020) resulting in rooted and corrected trees inferring the duplication, transfer and loss events that best reconcile the protein family trees with the species tree in terms of maximum likelihood. We used GeneRax with the following parameters: --rec-model UndatedDTL --prune-species-tree --max-spr-radius 3 -- reconciliation-samples 10. iTOL v4 (Letunic and Bork, 2019) was used to draw and annotate inferred protein trees reconciled with the species tree.

Reconciled phylogenetic trees in Newick format are provided in Supplementary Data 1-8. All proteins in the trees were annotated with species names, protein names and protein IDs; e.g., Hor.vu-DCL5|HORVU.MOREX.r3.1HG0044790.1 represents the DCL5 protein in barley (*Hordeum vulgare*). A complete list of all proteins included in the protein trees reported in this work are provided in Supplementary Table 2.

### Identifying Nuclear localization signals and catalytic triads of AGOs

NLS sequences were identified by scanning AGO protein sequences using LOCALIZER v1.0.5 (Sperschneider et al., 2017). To annotate catalytic residues located in the PIWI domain, multiple sequence alignments were performed on protein sequences of each AGO protein clade using MAFFT v7.505 (Katoh and Standley, 2013; Katoh et al., 2017; Rozewicki et al., 2019) with the following parameters: --anysymbol --dash -- originalseqonly --auto. Sequence alignments were first visualized using the SnapGene software (from Insightful Science; available at <u>snapgene.com</u>) to identify the coordinates of catalytic residues previously described (Qi et al., 2006; Carbonell et al., 2012; Zhang et al., 2014). Aligned residues matching the catalytic triad and conserved protein motifs were extracted using extractalign from EMBOSS v6.6.0.0 (Rice et al., 2000). Conserved protein motifs were visualized using WebLogo3 (Crooks et al., 2004).

## ACKNOWLEDGMENTS

We thank Danforth Center colleagues Elizabeth Kellogg and Yunqing Yu for helpful discussions, and Joanna Friesner for assistance with editing. This work was partly supported by a USDA National Institute of Food and Agriculture “BTT EAGER” award no. 2018–09058 (to B.C.M.). Sébastien Bélanger was supported by a postdoctoral fellowship from the Natural Sciences and Engineering Council of Canada (NSERC).

## AUTHOR CONTRIBUTIONS

SB, JZ, and BCM designed the analyses. SB performed the analyses. SB, JZ, and BCM interpreted the results. SB, JZ, and BCM wrote the manuscript.

## SUPPLEMENTAL FIGURES AND DATA

***Supplemental Figure 1.*** *Maximum-likelihood phylogeny of all AGO proteins annotated in the analyzed species. **(A)** Rooted phylogenetic tree with protein IDs of AGOs shown. **(A)** Unrooted view of the major clades of the phylogenetic tree in (A).*

***Supplemental Figure 2.*** *Rooted maximum-likelihood phylogeny of all SGS3 proteins annotated in the analyzed species.*

***Supplemental Figure 3.*** *Rooted maximum-likelihood phylogeny of all DRB proteins annotated in the analyzed species.*

***Supplemental Figure 4.*** *Rooted maximum-likelihood phylogeny of all SE proteins annotated in the analyzed species.*

***Supplemental Figure 5.*** *Rooted maximum-likelihood phylogeny of all HEN1 proteins annotated in the analyzed species.*

***Supplemental Table 1.*** *Sources of all proteomes analyzed in this study and their taxonomic classification.*

***Supplemental Table 2.*** *List of proteins annotated in the seven protein families and their phylogenetic-inferred protein names*.

***Supplemental Data 1.*** *Species tree of the 207 genomes analyzed in this study in Newick format*.

***Supplemental Data 2.*** *Maximum likelihood tree of 1,440 RDR proteins in Newick format*.

***Supplemental Data 3.*** *Maximum likelihood tree of 1,036 DCL proteins in Newick format*.

***Supplemental Data 4.*** *Maximum likelihood tree of 2,979 AGO proteins in Newick format*.

***Supplemental Data 5.*** *Maximum likelihood tree of 455 SGS3 proteins in Newick format*.

***Supplemental Data 6.*** *Maximum likelihood tree of 2,000 DRB proteins in Newick format*.

***Supplemental Data 7.*** *Maximum likelihood tree of 470 SE proteins in Newick format*.

***Supplemental Data 8.*** *Maximum likelihood tree of 224 HEN1 proteins in Newick format*.

