## Supplemental Figure 1 for "Phylogenetic analyses of AGO/DCL/RDR proteins in green plants refine the evolution of small RNA pathways"

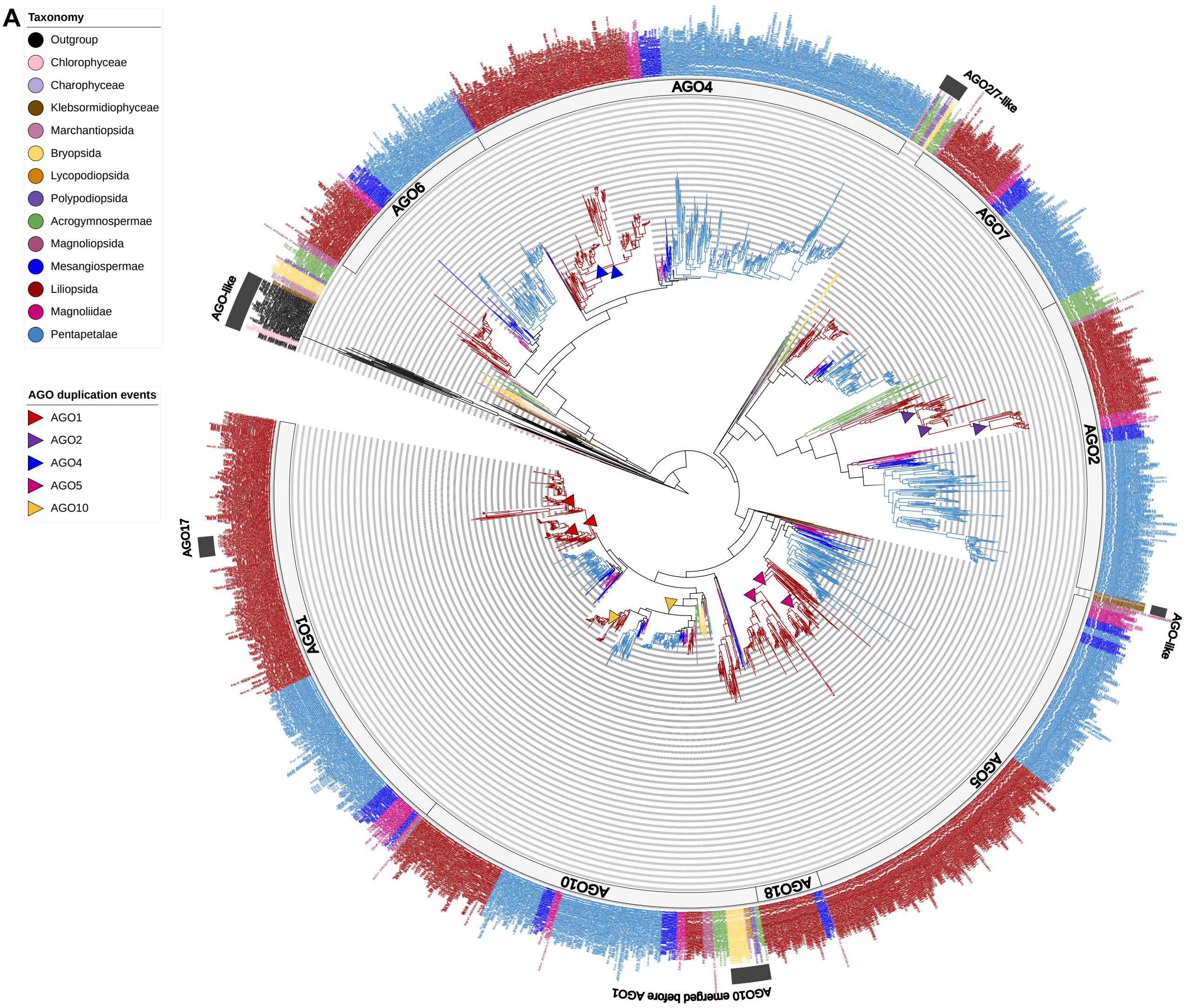


**2,979 AGO proteins**

(2,927 AGOs from Viridiplantae species)

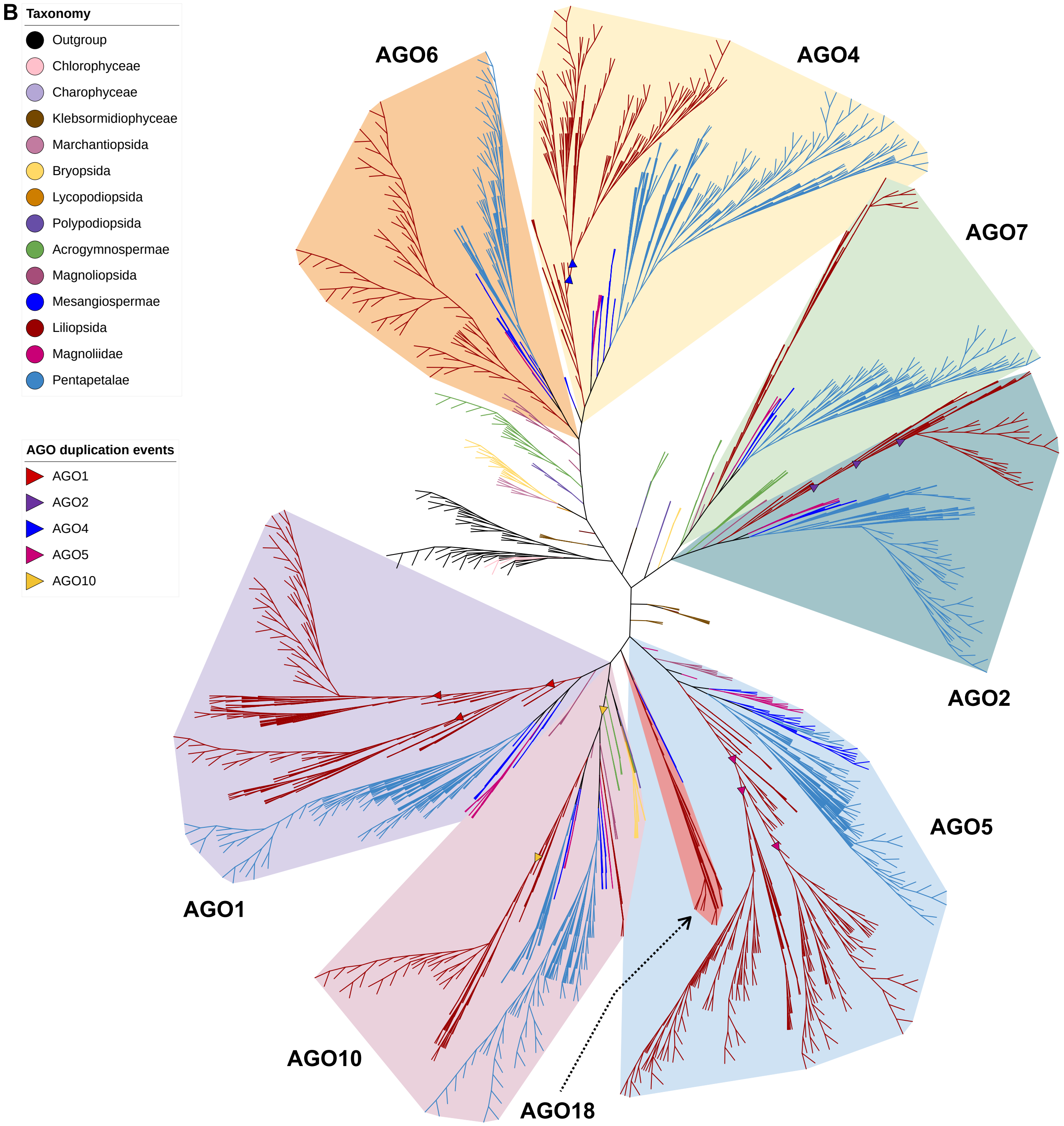


***Supplemental Figure 1.*** *Maximum-likelihood phylogeny of all AGO proteins annotated in the analyzed species.*
**(A)** Rooted phylogenetic tree with protein IDs of AGOs shown. **(B)** Unrooted view of the major clades of the phylogenetic tree in (A).
