## Supplemental Figure 2 for "Phylogenetic analyses of AGO/DCL/RDR proteins in green plants refine the evolution of small RNA pathways"

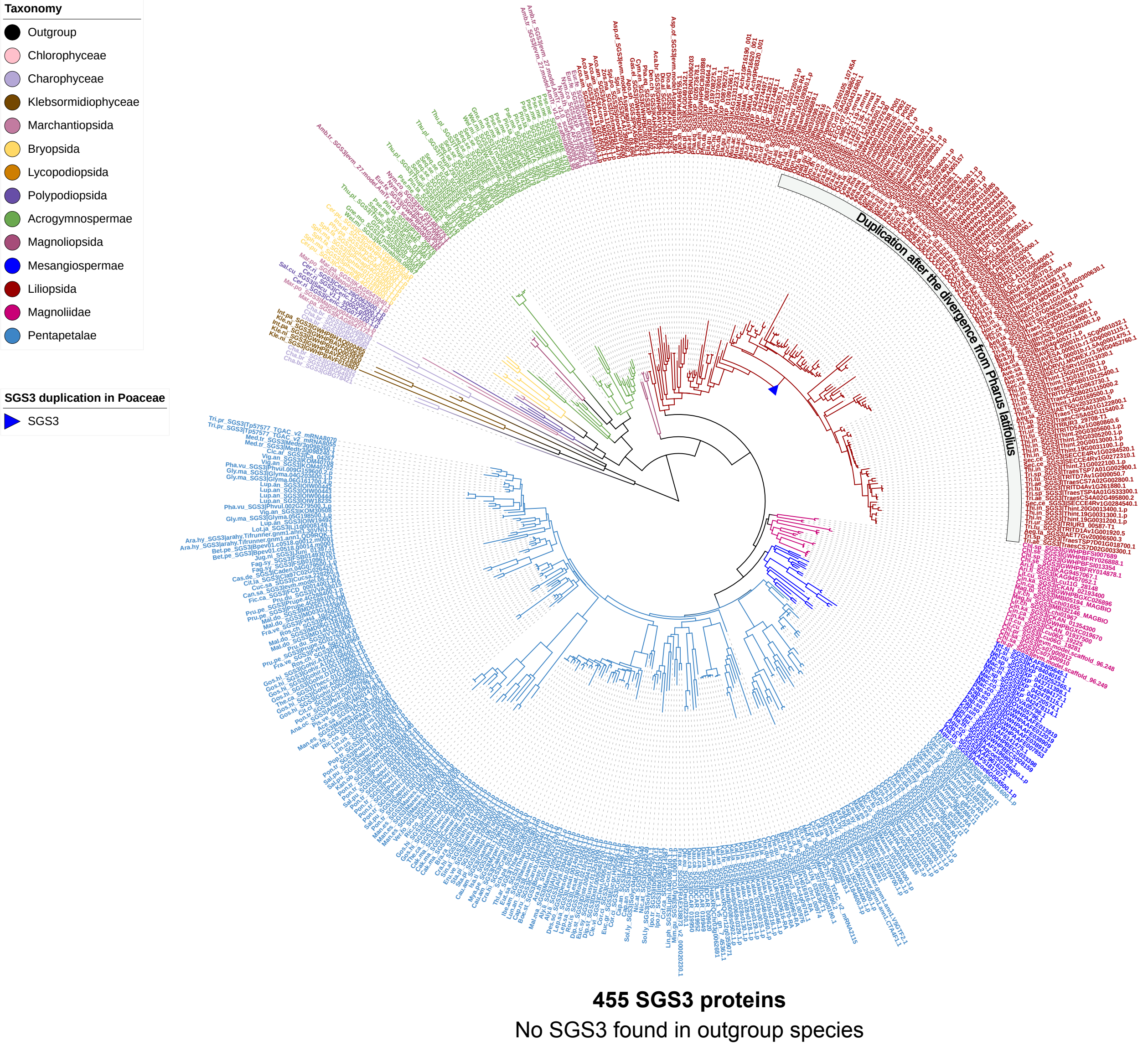


***Supplemental Figure 2.*** *Rooted maximum-likelihood phylogeny of all SGS3 proteins annotated in the analyzed species.*
