## Supplemental Figure 3 for "Phylogenetic analyses of AGO/DCL/RDR proteins in green plants refine the evolution of small RNA pathways"

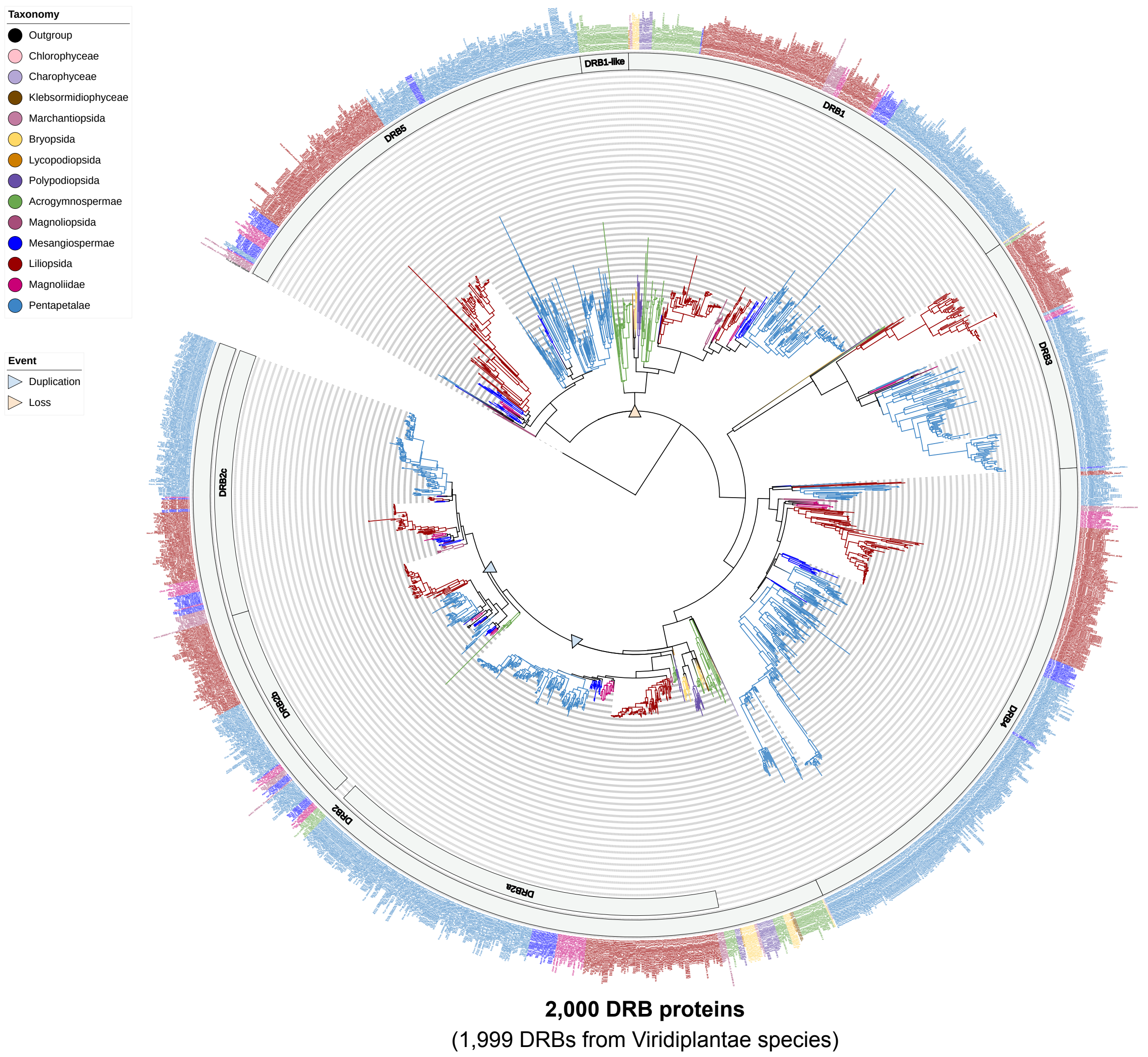


***Supplemental Figure 3.*** *Rooted maximum-likelihood phylogeny of all DRB proteins annotated in the analyzed species.*
