## Supplemental Figure 4 for "Phylogenetic analyses of AGO/DCL/RDR proteins in green plants refine the evolution of small RNA pathways"

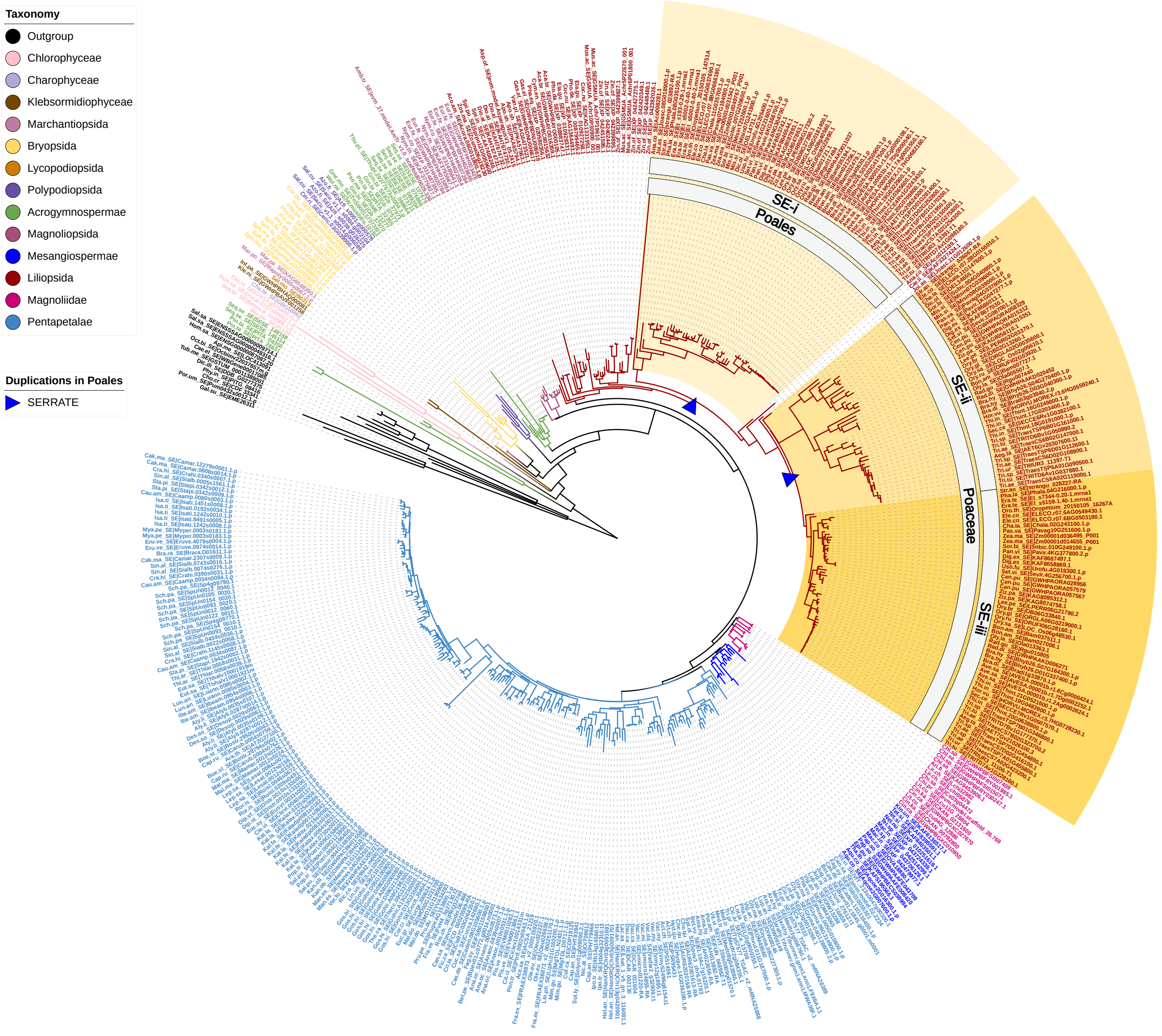


**470 SE proteins**

(458 SEs from Viridiplantae species)

***Supplemental Figure 4.*** *Rooted maximum-likelihood phylogeny of all SE proteins annotated in the analyzed species.*
