## Supplemental Figure 5 for "Phylogenetic analyses of AGO/DCL/RDR proteins in green plants refine the evolution of small RNA pathways"

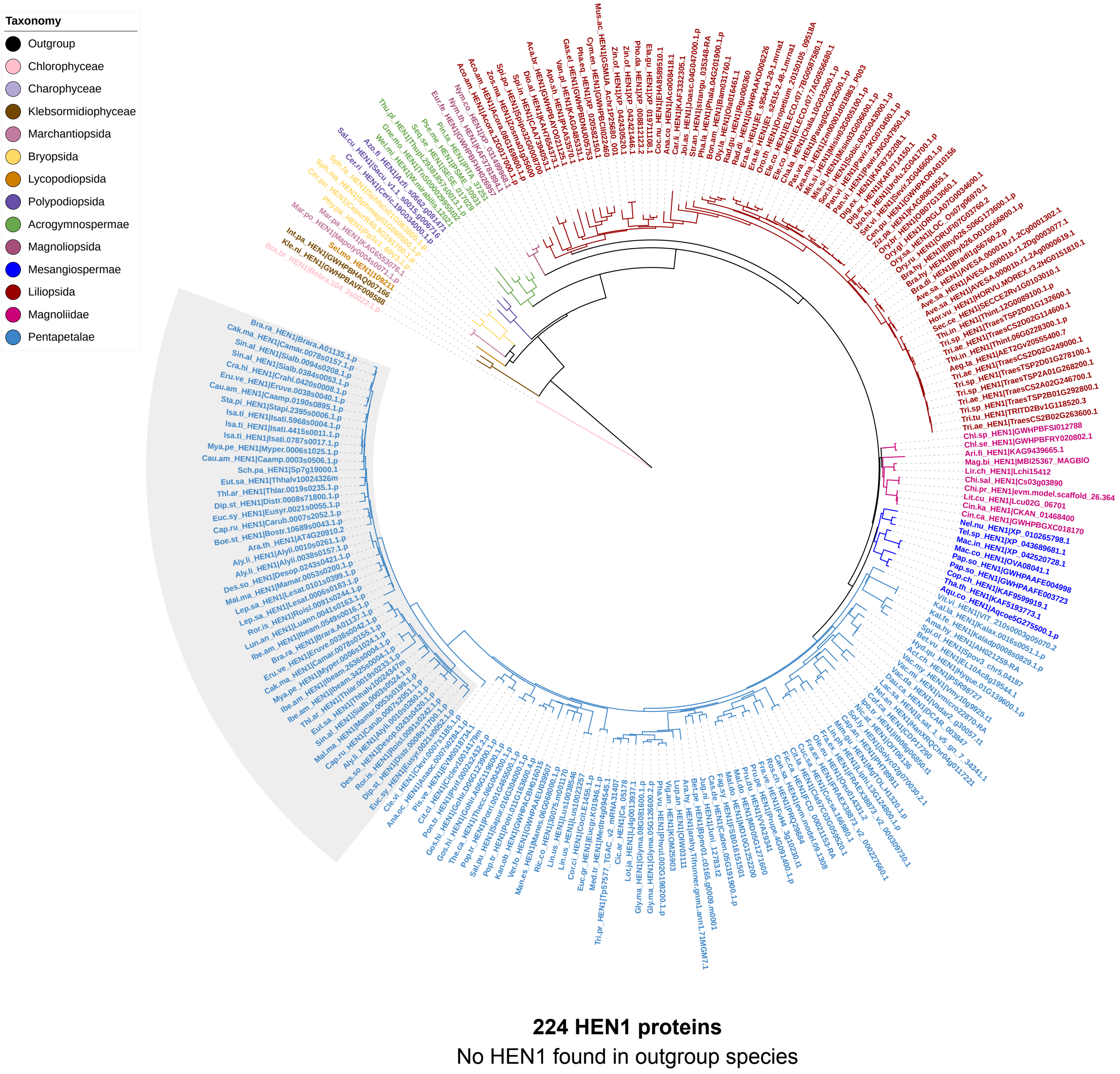


***Supplemental Figure 5.*** *Rooted maximum-likelihood phylogeny of all HEN1 proteins annotated in the analyzed species.*
